# Proteasome Regulation by Reversible Tyrosine Phosphorylation at the Membrane

**DOI:** 10.1101/2020.09.20.305532

**Authors:** Lu Chen, Xin Shu, Qiong Chen, Tiantian Wei, Xiaorong Wang, Qirou Wu, Heman Wang, Xiaoyan Liu, Xiaomei Zhang, Yanan Zhang, Suya Zheng, Lan Huang, Junyu Xiao, Chao Jiang, Zhiping Wang, Bing Yang, Xing Guo

## Abstract

Reversible phosphorylation has emerged as an important mechanism for regulating 26S proteasome function in health and disease. Over 100 phospho-tyrosine (pTyr) sites of the human proteasome have been detected, and yet their function and regulation remain poorly understood. Here we show that the 19S subunit Rpt2 is phosphorylated at Tyr439, a strictly conserved residue within the C-terminal HbYX motif of Rpt2 that is essential for 26S proteasome assembly. Unexpectedly, we found that Y439 phosphorylation depends on Rpt2 membrane localization mediated by its N-myristoylation. Multiple receptor tyrosine kinases (RTKs) can trigger Rpt2-Y439 phosphorylation by activating Src, a N-myristoylated tyrosine kinase. Src directly phosphorylates Rpt2-Y439 *in vitro* and negatively regulates 26S proteasome integrity and activity at cellular membranes, which can be reversed by the membrane-associated isoform of protein tyrosine phosphatase non-receptor type 2 (PTPN2). In H1975 lung cancer cells with activated Src, blocking Rpt2-Y439 phosphorylation by the Y439F mutation conferred partial resistance to the Src inhibitor saracatinib both *in vitro* and in a mouse xenograft tumor model, and caused significant changes of cellular responses to saracatinib at the proteome level. Our study has defined a novel mechanism involved in the spatial regulation of proteasome function and provided new insights into tyrosine kinase inhibitor-based anti-cancer therapies.

## Introduction

The 26S proteasome plays a central role in selective protein degradation in eukaryotic cells and its function is intricately regulated in health and disease (1–3). Consisting of at least 33 unique subunits, the 26S proteasome can be divided into two large subcomplexes, the 19S regulatory particle (RP) and the 20S core particle (CP). The 20S CP is formed by the stacking of four ring-shaped structures, each comprising seven α subunits (α1-7) or seven β subunits (β1-7), in an α-β-β-α order. Enclosed in the CP are the catalytic sites from the β1/2/5 subunits that are responsible for protein substrate hydrolysis. Substrate access to the catalytic core is usually restricted by N-terminal sequences of the α subunits that form a gate at the center of the α ring, which can be opened by binding of the 19S RP or other proteasome activators to the CP (4, 5).

The 19S RP contains six AAA^+^-type ATPase subunits (Rpt1-6) and thirteen non-ATPase subunits (Rpn1-3, 5-13 and 15). ATP binding and hydrolysis by the ATPases are crucial for substrate engagement, processing and translocation (6, 7). In addition, Rpt1-6 are essential for RP-CP association as they are assembled into a heterohexameric ring that directly docks on the α ring of CP. The C-terminal tails of Rpt1-6, which are highly conserved through evolution, can insert into cognate pockets formed by adjacent α subunits along the α ring. In particular, Rpt2, 3 and 5 have a signature HbYX motif (hydrophobic residue-tyrosine-any amino acid) at their very C-termini, featuring an invariant tyrosine residue at the penultimate position. Numerous genetic, biochemical and structural studies have demonstrated the critical importance of the Rpt tails for proteasome assembly, activity and cell viability (8–12).

Proteasome function is regulated at multiple levels to maintain cellular homeostasis (2, 13), while proteasome dysregulation has been implicated in a variety of human diseases such as neurodegenerative disorders and cancer (14–16). Reversible phosphorylation has emerged as an important mechanism for fine-tuning proteasome assembly, activity and subcellular localization (17, 18). Hundreds of phosphorylation sites have been found on the human 26S proteasome by mass spectrometry, nearly a third of which (143/453) are phospho-tyrosine (pTyr) sites (17). However, our understanding of the function and regulation of proteasome tyrosine phosphorylation has been extremely limited, except for studies on a couple of pTyr sites of the 20S subunit α4 (19, 20). A common theme from most phospho-proteomic investigations is that tyrosine phosphorylation of the proteasome mostly occurs in cancer cells with aberrant tyrosine kinase (TK) signaling (17).

Hyperactivated tyrosine phosphorylation often acts as a driving force behind tumorigenesis (21, 22). Inhibitors against receptor tyrosine kinases (RTKs, e.g. EGFR) and non-receptor tyrosine kinases downstream of the RTKs (such as Src and Abl) have been important components of targeted therapies against cancer (23, 24). However, even in the presence of activated TKs, phospho-tyrosine levels are usually kept low (< 1% of the whole phospho-proteome, 25), as many protein tyrosine phosphatases (PTPs) very efficiently remove these modifications (26, 27). Several PTPs have also been implicated in the oncogenic process, and these enzymes once considered “undruggable” can now be pharmacologically targeted, representing a new approach for anti-cancer treatment (28–30).

In this study, we focused on Src- and PTPN2-regulated reversible phosphorylation of the 19S subunit Rpt2. A shared feature between Src and Rpt2 is that they are both co-translationally modified by N-myristoylation, which usually targets proteins to the membrane (31). PTPN2, also known as T cell PTP (TC-PTP), has multiple splicing isoforms. The longest isoform (called TC48) contains a membrane-targeting region that is absent in the other isoforms (32). We found that Src and PTPN2 (TC-48) control the phosphorylation of membrane-tethered Rpt2 at Tyr439, which is the very tyrosine residue within the HbYX motif critical for 20S association. Src downregulates the abundance and activity of 26S proteasome at cellular membranes, while lung cancer cells expressing the non-phosphorylatable Rpt2-Y439F mutant are more resistant to Src inhibition *in vitro and in vivo*. These findings have revealed a new mechanism of spatial regulation of the proteasome, and suggest a considerable role of proteasome phosphorylation in determining the clinical outcome of tyrosine kinase inhibitors (TKIs).

## Results

### Rpt2-Y439 phosphorylation impairs 26S proteasome activity

According to the PhosphoSitePlus database (www.phosphosite.org), Rpt2-Y439 is the most frequently detected pTyr site of all 19S subunits. This is intriguing since Y439 is the exact tyrosine residue of the C-terminal HbYX motif of Rpt2 (L^438^<u>Y</u>^439^L^440^), which is strictly conserved among species (Fig. 1A). Consistent with the proteomics result, Y439 appeared to be the major tyrosine phosphorylation site of Rpt2 as the Y439F mutation largely diminished the total pTyr signal (Supplementary Fig. S1A). To confirm the occurrence of Rpt2-Y439 phosphorylation, we generated a phospho-specific antibody against this site. In 293T cells treated with pervanadate, a potent generic inhibitor of PTPs, Y439 phosphorylation could be detected from exogenously expressed Rpt2 with an internal Flag tag (“IF”, see below) following anti-Flag immunoprecipitation (Fig. 1B). Endogenous Rpt2 co-purified with TBHA-tagged Rpn1 subunit of the 19S RP (33, 34) was also found to be Y439-phosphorylated in the presence of pervanadate (Fig. 1C). Moreover, Y439 phosphorylation was evident from Rpt2 directly immunoprecipitated from rat embryonic forebrain tissues, although the phosphorylation was barely detectable in the midbrain from the same embryos (Fig. 1D). These initial characterizations indicate that Rpt2-Y439 phosphorylation does occur in cultured cells and *in vivo*, while its level may be dynamically regulated in different cell types or under different conditions (see also Supplementary Fig. S3C).

**Fig. 1.**
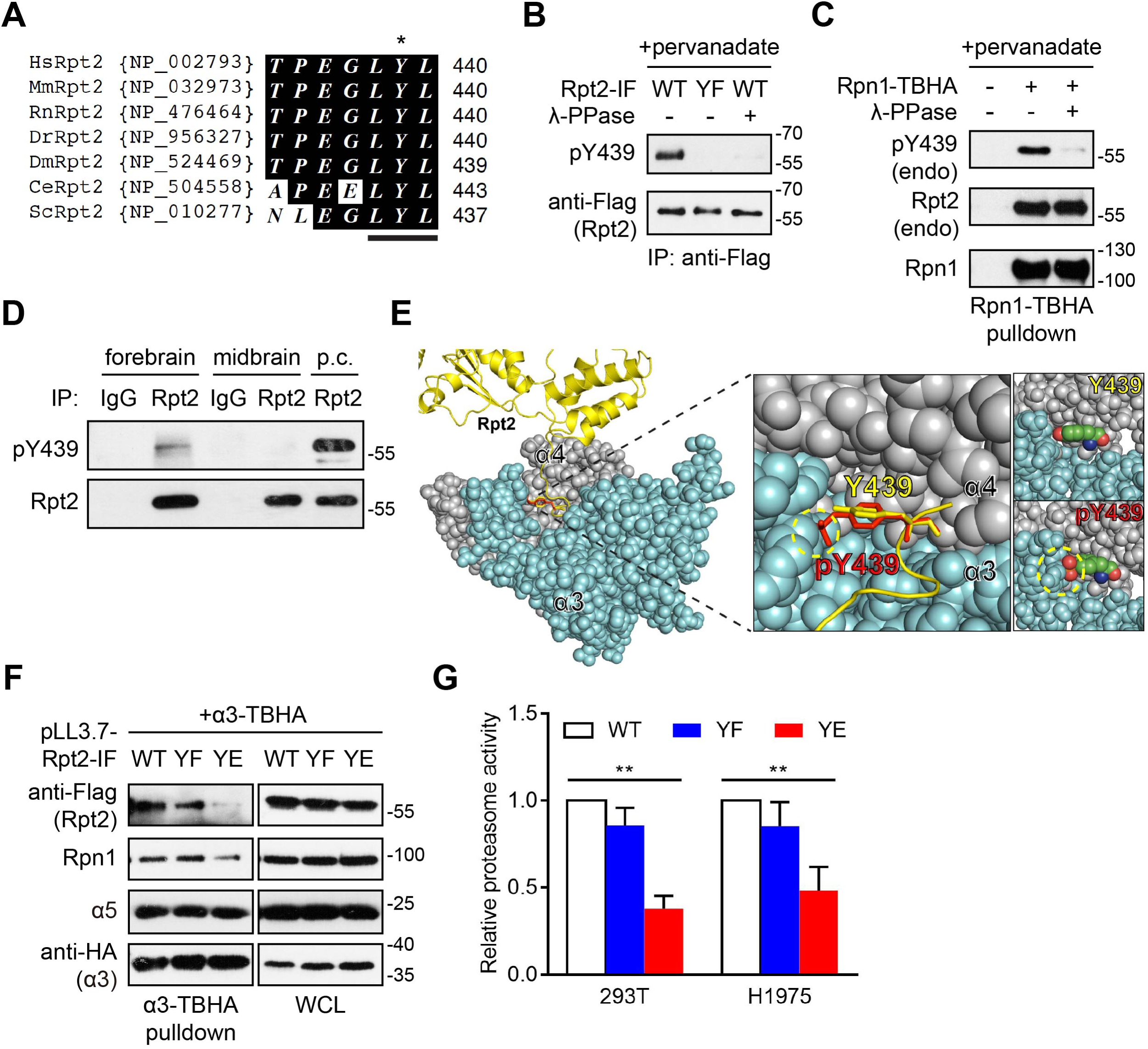
Rpt2-Y439 phosphorylation perturbs 26S proteasome function. **A.** Sequence alignment of the C-termini of Rpt2 from different species. The HbYX motif (LYL, underlined) including Y439 (*) is strictly conserved through evolution. **B.** 293T cells were transfected with the indicated variants of Rpt2-IF (internal Flag) and treated with pervanadate (0.1 mM, 15 min) before lysis. After anti-Flag immunoprecipitation, samples were treated with or without λ-phosphatase and probed with the indicated antibodies. YF, Y439F. **C.** 293T cells were transfected with Rpn1-TBHA, treated with pervanadate as in (B) and lysed for streptavidin pulldown. Samples were treated with or without λ-phosphatase and probed with the indicated antibodies. **D.** The indicated brain regions were dissected from E15.5 rat embryos. Tissue homogenates were incubated with a control rabbit IgG or anti-Rpt2 antibody for immunoprecipitation followed by western blot. p.c., positive control (pervanadate-treated cell extract). **E.** Structural modeling of Rpt2-Y439 phosphorylation. (Left and middle) Rpt2 is shown as yellow ribbons. Y439 and pY439 are modeled as yellow and red sticks, respectively. The dotted yellow circle indicates the steric clash between the phosphate group of pY439 and α3 (cyan spheres). (Right) Sphere presentation of Y439 and pY439 atoms, with O, C and N shown as red, green and blue, respectively. The model was built on the 26S proteasome structure 6MSJ (PDB ID) and further adjusted in Coot. **F.** 293T cells were stably transduced with the indicated pLL3.7 viruses for knockdown of endogenous Rpt2 and simultaneous expression of exogenous Rpt2-IF-WT/Y439F/Y439E. α3-TBHA was then transiently expressed in these cells for streptavidin pulldown and western blot analysis. WCL, whole cell lysate. **G.** 293T and H1975 cells transduced with the same pLL3.7 viruses as in (F) were lysed for proteasome activity measurement using Suc-LLVY-AMC as substrate. **, *P* < 0.01 (One-way ANOVA, n = 3). Paired comparisons: 293T WT vs. YF, *P* = 0.2012; 293T WT vs. YE, *P* = 0.0070. H1975 WT vs. YF, *P* = 0.3045; H1975 WT vs. YE, *P* = 0.0341.

From all previous studies and our structural modeling (Fig. 1E), it is immediately conceivable that Y439 phosphorylation should be incompatible with Rpt2-CP association due to a steric clash between the Rpt2 tail and the cognate pocket formed by α3 and α4. To examine the consequence of Rpt2-Y439 phosphorylation, we engineered stable cell lines in which endogenous Rpt2 was knocked down by shRNA and simultaneously replaced with WT or mutant Rpt2 (Supplementary Fig. S1B-E). When we used α3-TBHA to pull down the proteasome from these cells, the phosphomimetic mutant Rpt2-Y439E showed poor interaction with the 20S, while the phospho-deficient Rpt2-Y439F remained capable of binding the CP (Fig. 1F). Conversely, in Rpn11-TBHA pulldown assays, WT Rpt2 and both of the Y439 mutants appeared to be well incorporated into the 19S RP, but the amount of associated 20S was largely reduced in cells expressing Rpt2-Y439E (Supplementary Fig. S1C). Consistent with its defect in mediating RP-CP interaction, the Rpt2-Y439E mutant caused a significant inhibition of proteasome activity as compared to Rpt2-WT or Y439F (Fig. 1G). These results support the notion that Rpt2-Y439 phosphorylation may block RP-CP association and attenuate proteasome activity.

### Rpt2-Y439 phosphorylation occurs at the membrane

We then investigated how Rpt2-Y439 phosphorylation is regulated in cells and made a surprising discovery that Rpt2 tyrosine phosphorylation depends on its membrane localization. In our parallel study focusing on Rpt2 N-myristoylation, we engineered the expression construct of Rpt2-IF mentioned above, where aa. 85-91 of WT Rpt2 (MKPLEEK, an unstructured region, 35) was replaced by the Flag tag sequence, DYKDDDDK (Fig. 2A). This version of Rpt2 preserves its N-myristoylation, an absolutely conserved co-translational modification (31, 36–42) that is occluded in the commonly used Flag-Rpt2 because of N-terminal tagging. Rpt2-IF could be similarly incorporated into the 26S proteasome as endogenous Rpt2 and fully support proteasome functions, and was indistinguishable from the N-terminally tagged Flag-Rpt2 in their protein expression levels (Supplementary Fig. S2). Strikingly, however, Rpt2-IF showed much stronger overall tyrosine phosphorylation and Y439 phosphorylation than Flag-Rpt2 following pervanadate treatment (Fig. 2B). For this reason, Rpt2-IF was used instead of Flag-Rpt2 throughout this study.

**Fig. 2.**
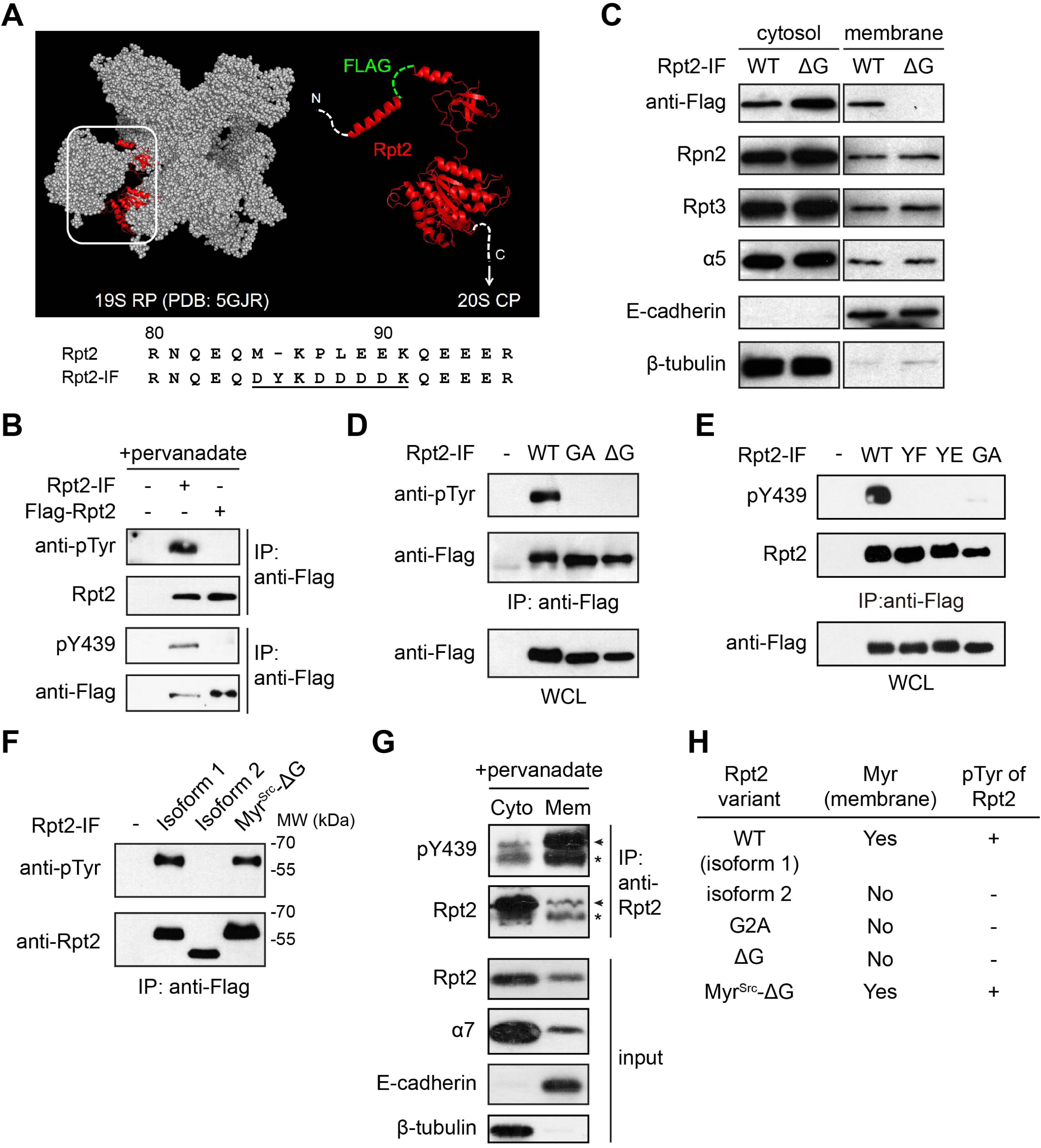
Rpt2 membrane localization is required for its tyrosine phosphorylation. **A.** In the cyro-EM structure of the 19S RP (grey spheres, top), Rpt2 (red ribbons) contains a flexible region (aa. 85-91, green dotted line), which was replaced with DYKDDDDK (Flag tag) to generate Rpt2-IF (bottom). **B.** Rpt2-IF and N-terminally tagged Flag-Rpt2 were transiently expressed in 293T cells. After pervanadate treatment, exogenous Rpt2 was immunoprecipitated and examined for total tyrosine phosphorylation or pY439 using the indicated antibodies. **C.** Rpt2-IF-WT and the myristoylation-deficient mutant Rpt2-IF-ΔG were ectopically expressed in 293T cells. Cytosolic and membrane fractions were separated and probed with the indicated antibodies. E-cadherin and β-tubulin served as membrane and cytosolic markers, respectively. **D.** The indicated Rpt2-IF constructs were transiently expressed in 293T cells. After pervanadate treatment, the Rpt2 variants were enriched by anti-Flag immunoprecipitation and probed for pTyr. GA, G2A. **E.** The indicated Rpt2-IF variants were expressed, immunoprecipitated as in (D) and blotted with anti-pY439 antibody. **F.** The indicated Rpt2-IF variants were expressed, immunoprecipitated and blotted as in (D). Isoform 1 is the same full-length Rpt2-IF-WT used elsewhere. Isoform 2 lacks aa. 1-73 of isoform 1. **G.** H1975 cells were treated with pervanadate and fractionated as in (C). Equal amounts of cytosolic and membrane fractions were immunoprecipitated with anti-Rpt2 antibody. pY439 and Rpt2 were probed (arrowheads). *, non-specific band. **H.** Summary of results from (D) to (F), highlighting the dependence of Rpt2 tyrosine phosphorylation on its membrane localization.

The proteasome has been found to associate with different membranes of the cell (43, 44). Both CP and RP subunits could be detected in the membrane fractionation of cells by western blot (Fig. 2C). Ectopically expressed Rpt2-IF mutants (38) with the myristoylation site Gly2 deleted (ΔG) or mutated (G2A) lost their membrane localization and tyrosine phosphorylation (including pY439) simultaneously (Fig. 2C-E). The same was true for a putative splicing variant of Rpt2 (NP_001317141) missing the N-terminal 73 amino acids of the full-length protein (Fig. 2F). On the contrary, tyrosine phosphorylation of Rpt2-IF-ΔG was fully restored by fusing the commonly used myristoylation sequence of Src (45) to its N-terminus (Fig. 2F). Cell fractionation assays further revealed that the Rpt2-pY439 signal was predominantly found in the membrane fraction (Fig. 2G). These results demonstrate that Rpt2-Y439 phosphorylation depends on its N-terminal myristoylation and occurs primarily at membrane structures of the cell (Fig. 2H).

### Src phosphorylates Rpt2-Y439

Next, we set out to search for the Rpt2-Y439 kinase(s), which is likely to be a membrane-localized protein. An important hint came from a previous phosphoproteomic study that the EGFR inhibitor gefitinib (Iressa^®^) completely suppressed Rpt2-Y439 phosphorylation in H3255 non-small cell lung cancer (NSCLC) cells without affecting other phospho-tyrosine sites of the proteasome (46). We confirmed this phenomenon by western blot (Supplementary Fig. S3A), and observed a similar inhibition of pY439 by the third-generation EGFR inhibitor, osimertinib (AZD9291, Tagrisso^®^), in another NSCLC line H1975 that bears the L858R/T790M mutations of EGFR (47) (Supplementary Fig. S3B). Consistently, Y439 phosphorylation of endogenous Rpt2 could be enhanced by EGF in both 293T and H1975 cells (Supplementary Fig. S3C), with the basal level of pY439 being higher in the latter likely due to the activating EGFR mutations. In addition to EGFR, overexpression of several other RTKs such as PDGFRβ, FGFR1, FGFR3, RET and NTRK1 all markedly enhanced Rpt2-Y439 phosphorylation (Fig. 3A). Induction of pY439 is therefore not unique to EGFR but is likely to be mediated by a common effector downstream various RTKs. Interestingly, RTK-stimulated Y439 phosphorylation was completely blocked by Src inhibitors such as saracatinib/AZD0530 or PP1 (Fig. 3A). In addition, pervanadate-induced pY439 was also fully suppressed by Src inhibitors saracatinib, PP1 or dasatinib (Sprycel^®^), but not the Abl kinase inhibitor imatinib (Gleevec^®^) (Fig. 3B). This suggests that Src and/or Src family kinases (SFKs) are responsible for Rpt2-Y439 phosphorylation.

**Fig. 3.**
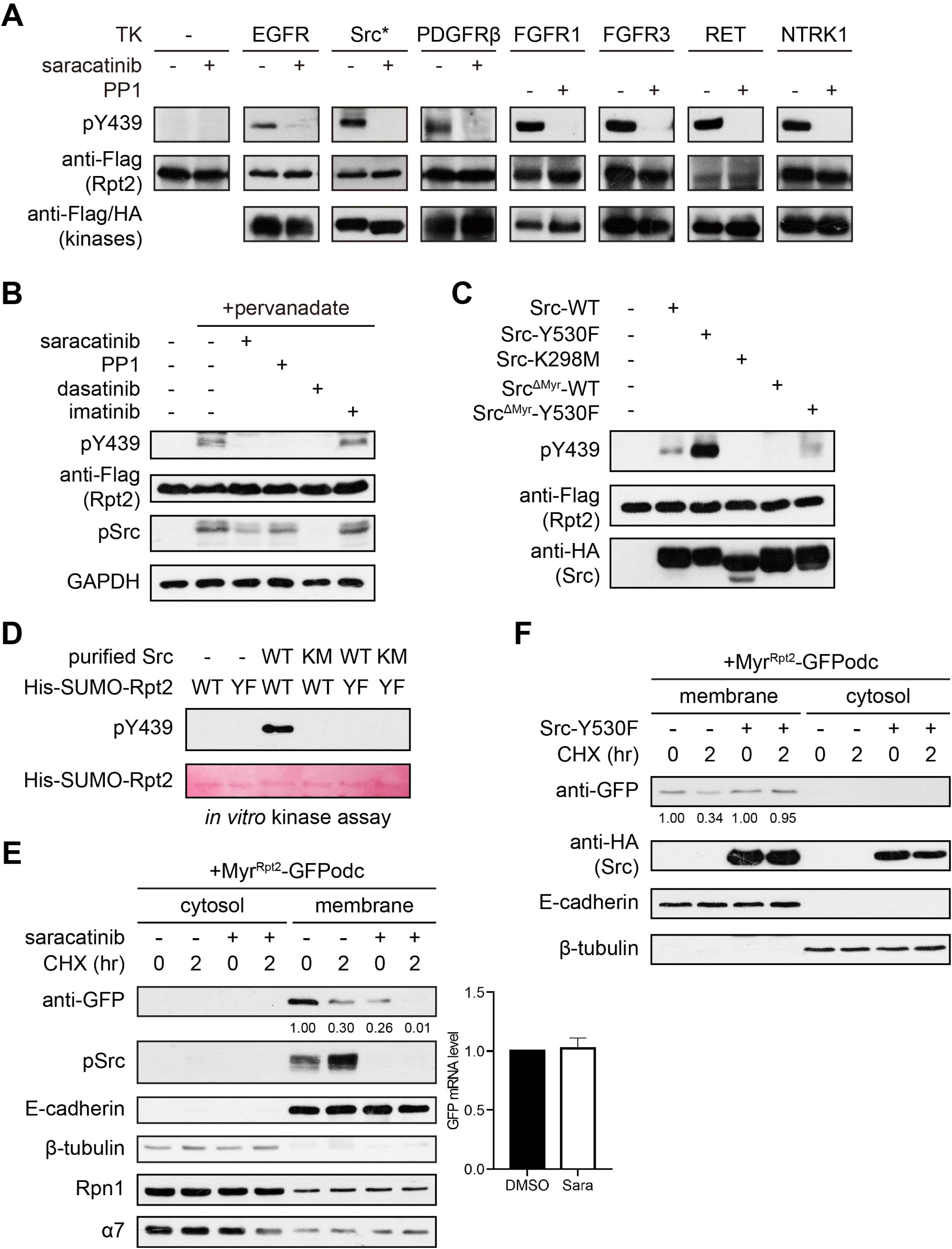
Src phosphorylates Rpt2-Y439. **A.** 293T pLL3.7-Rpt2-IF-WT cells transfected with vector control or the indicated tyrosine kinases (TKs) were treated with DMSO (-) or the indicated Src inhibitors (5 µM, 1 hr) before harvest. Whole cell lysates were probed with the antibodies shown. Src*, Src-Y530F-TBHA. All other TKs contain a C-terminal 3xFlag-V5 tag. **B.** 293T pLL3.7-Rpt2-IF-WT cells were treated with DMSO (-) or different inhibitors (5 µM, 1 hr). Pervanadate was added at 15 min before harvest. Whole cell lysates were probed with the indicated antibodies. pSrc, Src-pY419. **C.** 293T pLL3.7-Rpt2-IF-WT cells were transfected with vector control or variants of Src-TBHA. Whole cell lysates were probed with the indicated antibodies. **D.** Purified Src (WT or K298M) was incubated with recombinant His-SUMO-Rpt2 protein (WT or Y439F) for an *in vitro* kinase assay. Y439 phosphorylation was examined by western blot, and equal loading of His-SUMO-Rpt2 was shown by Ponceau S staining of the membrane. **E.** A single clone of 293T cells stably expressing Myr^Rpt2^-GFPodc were treated with DMSO or saracatinib (5 µM, 4 hrs). Before harvest, cells were treated with cycloheximide (CHX, 50 µg/ml) for the indicated time. After cell fractionation, the indicated proteins were examined by western blot (left). Relative levels of Myr^Rpt2^-GFPodc (normalized to the untreated sample) were quantified and shown. The mRNA levels of Myr^Rpt2^-GFPodc from DMSO and saracatinib-treated cells were determined by qPCR (right). **F.** The same reporter cell line as in (E) was transfected with vector or Src-Y530F-TBHA and. CHX was added and cells were processed as in (E).

We focused on Src, the primary target of saracatinib (48). Indeed, ectopic expression of wild-type Src and its constitutively active form Src-Y530F in 293T cells greatly increased Rpt2-Y439 phosphorylation, while the kinase-deficient mutant Src-K298M failed to do so (Fig. 3C). Src could co-purify with the proteasome from cells (Supplementary Fig. S3D) and directly phosphorylate recombinant Rpt2 at Y439 *in vitro* (Fig. 3D and Supplementary Fig. S3E). Notably, Src itself is also membrane-anchored via N-myristoylation (49). As expected, a Src mutant lacking its N-terminal myristoylation sequence (Src^ΔMyr^) was much less active toward Rpt2-Y439 than the wild-type kinase (Fig. 3C). These results provided an explanation for the dependence of Y439 phosphorylation on Rpt2 membrane localization. To further assess the activity of membrane-associated proteasomes, we generated a membrane-targeted version of the commonly used proteasome activity reporter GFPodc (50). This new reporter, designated Myr^Rpt2^-GFPodc, contains the N-terminal myristoylation sequence of Rpt2 (aa. 1-24) and could be successfully targeted to the membrane and degraded by the proteasome (Supplementary Fig. S3F). Using a single clone of 293T cells stably expressing this reporter, we found that Src inhibition by saracatinib not only reduced the steady-state level of Myr^Rpt2^-GFPodc but also greatly accelerated its protein degradation, without affecting its transcriptional expression (Fig. 3E). On the contrary, overexpression of activated Src essentially blocked Myr^Rpt2^-GFPodc degradation (Fig. 3F). These data have identified Src as a novel proteasome kinase that phosphorylates Rpt2-Y439 and regulates proteasome activity at the membrane.

### PTPN2 dephosphorylates Rpt2-pY439

Previously we performed Rpn11-HTBH pulldown and identified PTPN2/TC-PTP as a proteasome-interacting protein by mass spectrometry. Three other proteomic studies also documented such interaction using PTPN2 as bait (51-53, Supplementary Fig. S4A). Endogenous PTPN2 was readily co-purified with the proteasome by Rpn11-TBHA pulldown (Supplementary Fig. S4B). Reciprocally, anti-Flag-PTPN2 immunoprecipitation showed that endogenous proteasome subunits preferentially interacted with the membrane-associated isoform of PTPN2 (TC48) but exhibited little binding with the nuclear form, TC45 (Fig. 4A). Fluorescence microscopy studies also confirmed co-localization between Rpt2-IF and GFP-TC48 in cells (Supplementary Fig. S4C). These results prompted us to test whether PTPN2 regulates Rpt2 phosphorylation.

**Fig. 4.**
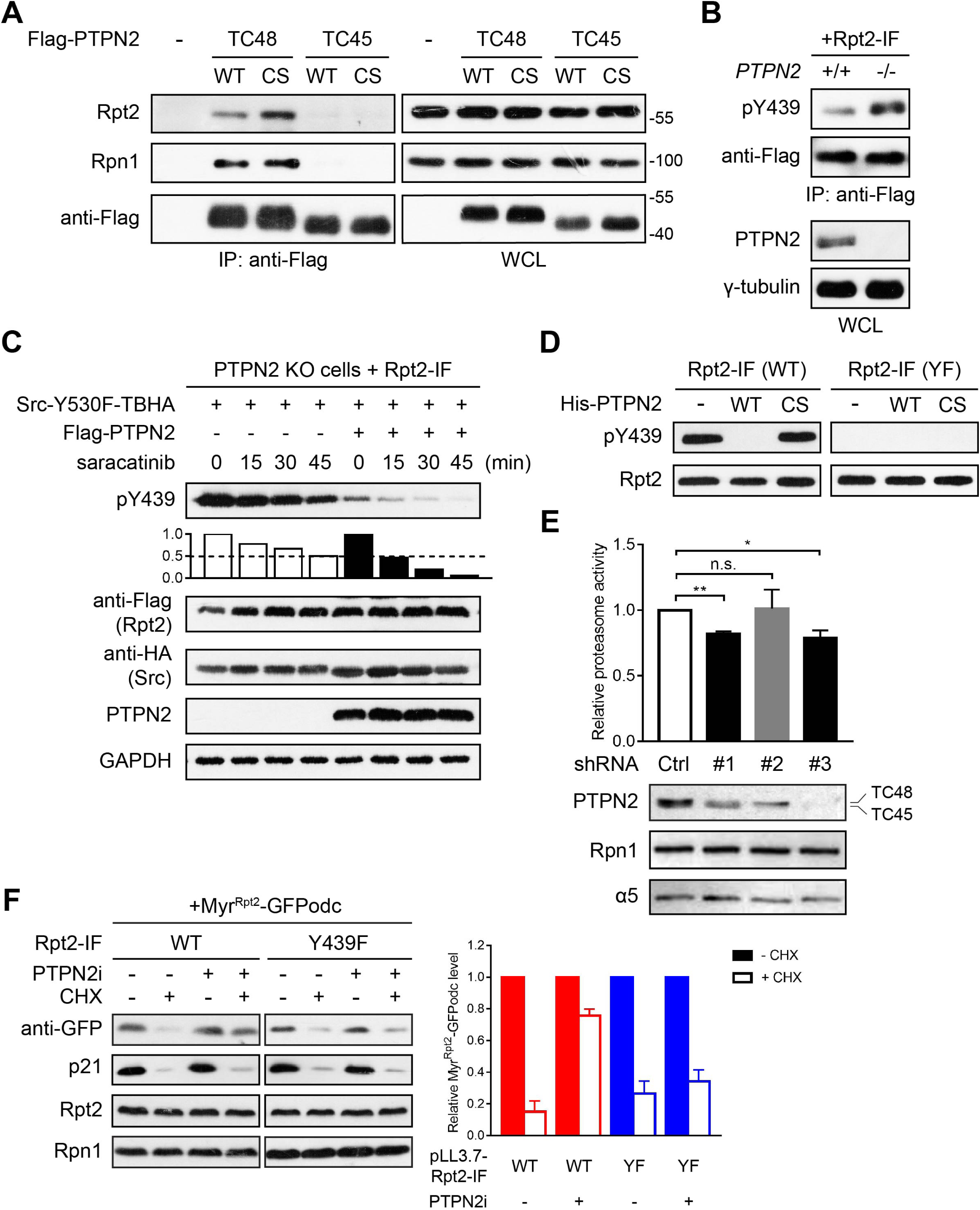
Rpt2-pY439 is dephosphorylated by PTPN2. **A.** Flag-PTPN2 isoforms were expressed in 293T cells. Co-immunoprecipitation of PTPN2 and endogenous proteasome subunits was determined by western blot. CS, C216S. **B.** Rpt2-IF was expressed in parental (+/+) or PTPN2-null (-/-) HeLa cells. After anti-Flag immunoprecipitation, pY439 was probed. **C.** PTPN2-null 293T cells were co-transfected with Rpt2-IF and activated Src plus vector control or Flag-PTPN2 (TC48). Saracatinib (100 nM) was added at the indicated time points before cell lysis. Whole cell lysates were probed with the indicated antibodies, and normalized levels of pY439 in each group (against time “0”) are shown. Open bars, vector control. Filled bars, TC48-transfected cells. **D.** Rpt2-IF-WT/Y439F was immunoprecipitated from pervanadate-treated 293T cells and incubated with bacterially purified His-PTPN2-WT/C216S in an *in vitro* phosphatase assay. Samples were blotted with the indicated antibodies. **E.** 293T cells were transfected with a scrambled control shRNA or three independent shRNAs targeting PTPN2. shRNA #1 specifically targets the 3’-UTR of TC48, #2 specifically targets the 3’-UTR of TC45, and #3 targets the coding sequence shared by the two isoforms. Whole cell lysates were used for proteasome activity measurement using Suc-LLVY-AMC as substrate. Knockdown efficiency was verified by the anti-PTPN2 blot. Rpn1 and α5 served as loading control. **, *P* < 0.01; *, *P* < 0.05; n.s., not significant (Student’s *t* test vs. control, n = 3). **F.** The same Myr^Rpt2^-GFPodc reporter line as in Fig. 3 was stably transduced with pLL3.7-Rpt2-IF-WT or Y439F. Cells were pretreated with PTPN2 inhibitor (Compound 8, 50 nM) or vehicle (ethanol) for 1 hr before CHX treatment (3 hrs). Whole cell lysates were probed with the indicated antibodies. Representative blots are shown on the left, and normalized Myr^Rpt2^-GFPodc levels from three independent experiments are shown on the right (filled bars, no CHX; open bars, with CHX).

CRISPR/Cas9-mediated PTPN2 knockout led to an elevation of Rpt2-pY439 in cells (Fig. 4B). Since PTPN2 dephosphorylates and inactivates Src (54), the increase of pY439 might have resulted from reduced dephosphorylation or elevated Src activity or both. To further dissect the role of PTPN2 in Rpt2 regulation, we co-expressed Rpt2 with activated Src to achieve a high starting level of pY439, then inhibited Src activity by saracatinib and monitored the rate of pY439 disappearance afterwards. In PTPN2-null cells, Src induced strong Rpt2-Y439 phosphorylation that gradually decreased with the addition of saracatinib, while re-expression of TC48 markedly accelerated the decline of pY439 (Fig. 4C). This result supports a direct role of PTPN2/TC48 in controlling Rpt2-pY439 in cells. Evidence from *in vitro* phosphatase assays using bacterially purified PTPN2 further demonstrated that Rpt2 is a direct substrate of PTPN2 (Fig. 4D and Supplementary Fig. S4D). In agreement with being a Rpt2-pY439 phosphatase, TC48 knockdown decreased proteasome peptidase activity, while depletion of TC45 had little affect (Fig. 4E). Furthermore, treating cells with a specific PTPN2 inhibitor, Compound 8 (55), impeded Myr^Rpt2^-GFPodc clearance in Rpt2-WT cells, while such effect of PTPN2 inhibition was lost upon replacing endogenous Rpt2 with Rpt2-Y439F (Fig. 4F). Degradation of the non-membrane substrate of the proteasome, p21^Cip1^, was not altered by the same treatment (Fig. 4F). Therefore, Rpt2-pY439 appears to be a major target through which PTPN2 positively regulates proteasome activity.

### Rpt2-Y439F mutation lowers cell sensitivity to saracatinib

To further explore the functional importance of Y439 phosphorylation, we turned to the aforementioned H1975 NSCLC cells, in which Src is active due to the EGFR mutations and Rpt2-Y439 phosphorylation was effectively inhibited by saracatinib (Supplementary Fig. S5A). Rpt2-IF-WT or Y439F were stably introduced using the same knockdown-addback approach and were expressed at similar levels comparable to that of endogenous Rpt2 in H1975 parental cells (Supplementary Fig. S1D, E). Saracatinib treatment clearly suppressed the viability of H1975-WT cells in colony formation assays in a dose-dependent manner (Fig. 5A). However, although H1975-YF cells were also growth-inhibited, they were apparently less sensitive to saracatinib than WT cells (Fig. 5A). A brief survey of several proliferation/survival-related proteins showed that p53 induction by saracatinib was present in both cell lines, whereas the level of cleaved PARP1 was higher in H1975-WT cells, indicating increased apoptosis (Fig. 5B). On the other hand, CDK4/cyclin D1 were maintained at higher levels in H1975-YF cells after saracatinib treatment, and CDK2 activation was also more evident in the YF cells probably due to elevated expression of cyclin H (Fig. 5B). These differences may partly explain the reduced sensitivity of H1975-YF cells to saracatinib, although the intracellular localizations of these proteins suggest that they may not be directly affected by saracatinib-mediated inhibition of Rpt2-Y439 phosphorylation at the membrane.

**Fig. 5.**
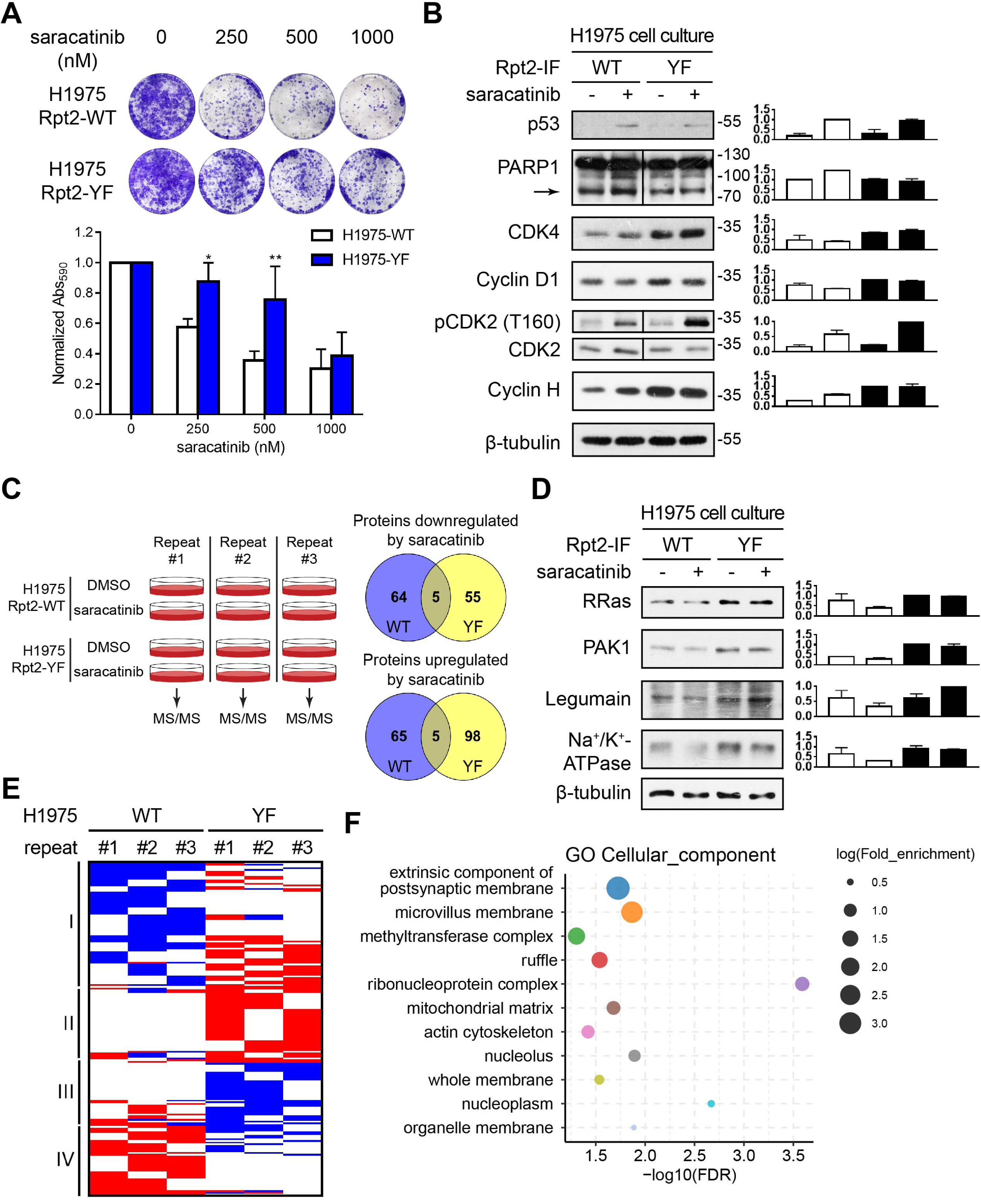
Altered responses to saracatinib in Rpt2-Y439F cells. **A.** Colony formation assays of H1975 cells transduced with pLL3.7-Rpt2-IF-WT or Y439F. Crystal violet-stained cells were photographed (top) and quantified (bottom). *, *P* < 0.05; **, *P* < 0.01 (unpaired two-tailed Student’s *t*-test, WT vs. YF, n = 3). **B.** H1975-WT/YF cells were treated with DMSO or saracatinib (250 nM) for 24 hrs. The indicated endogenous proteins were probed from whole cell extracts. Representative blots are shown on the left. Arrow, cleaved PARP1. Results from 3 independent blots of each protein were quantified and shown on the right. **C.** Left: The experimental setup for proteomic studies of H1975-WT/YF cells treated with DMSO or saracatinib (250 nM, 24 hrs). Right: Venn diagrams showing proteins that were downregulated (top) and upregulated (bottom) from each cell line in at least 2 of the 3 biological repeats. **D.** Western blot analysis of representative membrane-associated proteins differentially affected by saracatinib in H1975-WT and YF cells. Results from 3 independent blots of each protein were quantified and shown on the right. **E.** Heat map visualization of the 191 proteins that responded differently to saracatinib in H1975-WT and YF cells as revealed by mass spectrometry. Each protein in each repeat is represented by a horizontal line. Blue, downregulated by saracatinib (D = −1); red, upregulated by saracatinib (D = 1); white, unchanged by saracatinib (D = 0). **F.** Gene Ontology (GO) term analysis showing cellular components enriched among the 191 differentially regulated proteins. Only first-level terms in each hierarchy (indicated by a different color) with FDR < 0.05 are shown.

To gain further insight into the specific involvement of Rpt2-Y439 phosphorylation in mediating cellular responses to saracatinib, we compared the whole proteomes of H1975-WT and YF cells with or without saracatinib treatment using label-free mass spectrometry (Fig. 5C). The drug concentration (250 nM, or 135 ng/ml) was within the range of plasma concentrations in patients receiving saracatinib treatment in clinical trials (56). Of all identified proteins, 3036 proteins overlapped among all three biological repeats (Table S3). In every repeat, saracatinib caused upregulation of around 200∼300 proteins in each cell line (> 2-fold change in intensity), and a similar number of proteins were downregulated (< 0.5-fold) (Table S4). We assigned a D-score of +1, 0 or −1 to each upregulated, unchanged or downregulated protein, respectively, in every experiment. For any individual protein, its overall response to saracatinib in each cell line could be qualitatively determined by the sum of its D-scores (D_sum_) from the three repeats. There were 280 proteins with D_sum_ ≥ 2 or ≤ −2 in WT and YF cells altogether, such that they were upregulated or downregulated by saracatinib in at least two of the three runs (Table S4). Among them, 69 proteins consistently decreased in WT cells in response to saracatinib, the majority of which (41/69, 60%) are annotated as membrane-localized according to the UniProt database. However, most of these proteins (64/69) were not downregulated in the YF cells, and *vice versa* (Fig. 5C and Table S5). Saracatinib-upregulated proteins also minimally overlapped between WT and YF cells (Fig. 5C and Table S6). Western blot analysis of RRas, PAK1 and Legumain confirmed the mass spectrometry results that their protein levels decreased in saracatinib-treated H1975-WT cells but not in H1975-YF cells (Fig. 5D). A similar trend was seen with Na^+^/K^+^-ATPase (Fig. 5D). These proteins have been regarded as pro-tumorigenic and/or anti-cancer drug targets (57–60). Treating cells with the proteasome inhibitor bortezomib caused accumulation of RRas, PAK1 and Na^+^/K^+^-ATPase in the membrane fraction, suggesting that these proteins could be substrates of membrane-associated proteasomes (Supplementary Fig. S5B). In contrast, Legumain, which locates within the lysosome, was apparently not stabilized by bortezomib (Supplementary Fig. S5B). Therefore, its expression patterns in WT and YF cells might have resulted from other (indirect) mechanisms. Nonetheless, the Y439F mutation clearly had a profound impact on proteomic changes induced by saracatinib in H1975 cells.

Among the 280 up-/down-regulated proteins, 191 of them showed different responses to saracatinib between the two cell lines (defined as |D_sum (WT)_ - D_sum (YF)_| ≥ 2, Fig. 5E and Table S7). Gene Ontology (GO) analysis of these proteins again showed that they are enriched in membrane structures of the cell (Fig. 5F) and involved in various biological processes including vesicle trafficking, intracellular protein transport, organelle function/organization, cell growth and stress response (Supplementary Fig. S5C, D). These 191 proteins could be assigned to 4 groups (I – IV) based on their expression patterns following saracatinib treatment (Fig. 5E). Of particular interest were proteins that were selectively downregulated (Group I) or less upregulated in WT cells (Group II), some of which (including RRas and PAK1) could have undergone enhanced proteasomal degradation following saracatinib treatment in WT but not YF cells. Importantly, our literature search showed that approximately two thirds of them have been implicated in cancer, with the vast majority of them being pro-survival and/or oncogenic. Their downregulation in WT cells is in agreement with the cytotoxic activity of saracatinib, whereas the weakened or opposite changes of these proteins in the YF cells may underlie the elevated resistance to saracatinib as seen above.

### H1975-YF tumors gain partial resistance to saracatinib

Saracatinib is currently in Phase 2/3 clinical trials against several types of solid tumors (https://www.clinicaltrials.gov) and has been shown to suppress H1975 tumor formation in nude mice (61). We used a similar xenograft model to examine the relevance of Rpt2-Y439 phosphorylation to the anti-cancer function of saracatinib *in vivo*. H1975-WT and YF cells injected subcutaneously into nude mice formed palpable tumors at similar rates. Both groups of mice were then randomly divided to receive vehicle (DMSO) or saracatinib (25 mg/kg/d) treatment for 12 consecutive days (Fig. 6A). As reported (61), orally administered saracatinib inhibited Src activation in tumor cells (Fig. 6B) and reduced tumor growth in mice (Fig. 6C, D). However, the tumorigenic growth of H1975-YF cells was less suppressed by saracatinib than that of the H1975-WT cells (Fig. 6C, D), echoing results from the colony formation assays (Fig. 5A). Western blot analysis of tumor homogenates again revealed different ramifications of drug treatment in WT and YF tumors. Saracatinib caused a decrease of CDK4, RRas, Legumain and the cell surface molecule CD46/MCP in WT tumors but not in the YF tumors (Fig. 6E), consistent with the mass spectrometry study using cultured cells. Pro-tumorigenic kinases CDK2, Erk1/2 and PAK1 were also more active or present at higher amounts in YF tumors than in WT tumors after saracatinib treatment (Fig. 6E). These differences might at least in part account for the dampened effect of saracatinib on the growth of YF tumors. Taken together, these *in vivo* data further underscore the importance of Rpt2-Y439 phosphorylation in determining cellular sensitivity and responses to saracatinib (Fig. 6F). This form of proteasome regulation may thus be a clinically relevant mechanism that can be tapped to improve the outcome of TKI-based anti-cancer therapies.

**Fig. 6.**
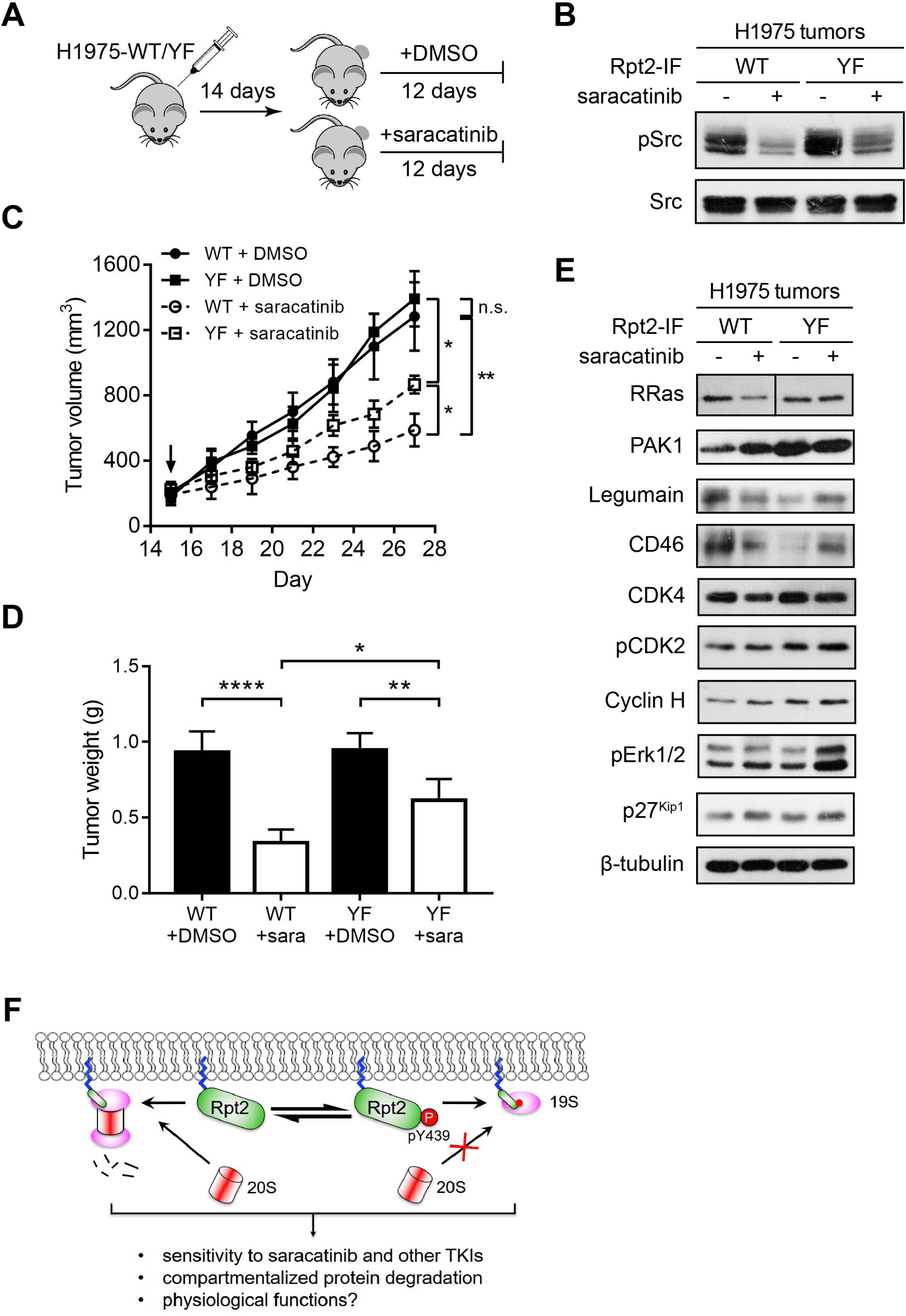
H1975-Y439F tumors were less inhibited by saracatinib. **A.** Schematic of mouse xenograft study with H1975-WT/YF cells. **B.** Tumors were dissected at the end of the study, homogenized and probed with the indicated antibodies to demonstrate effective Src inhibition by saracatinib treatment. **C.** Growth curves of tumors from the start of DMSO/saracatinib treatment (downward arrow). *, *P* < 0.05; **, *P* < 0.01 (unpaired two-tailed Student’s *t*-test between the indicated groups, n = 5). n.s., not significant. **D.** Tumor weight measurement. *, *P* < 0.05; **, *P* < 0.01; ****, *P* < 0.0001 (unpaired two-tailed Student’s *t*-test between the indicated groups, n = 5). **E.** Tumors were processed as in (B) and extracts were blotted with the indicated antibodies. **F.** Y439 phosphorylation of membrane-tethered Rpt2 affects 26S proteasome activity at the membrane and cellular responses to Src inhibition by saracatinib.

## Discussion

Proteasome tyrosine phosphorylation has been known for a long time (62) and constitutes a surprisingly high proportion of the proteasome phosphoproteome, with 9 of the top 10 most frequently detected proteasome phosphosites being pTyr (17). However, we noticed that in unstimulated cells such as 293T, detection of proteasome tyrosine phosphorylation is usually challenging. This discrepancy may be largely ascribed to differences between the detection methods, i.e. immunoblotting vs. high-sensitivity mass spectrometry aided by pervanadate treatment and phospho-peptide enrichment, but also raises questions about the general importance and regulatory mechanisms of proteasome tyrosine phosphorylation (19, 20, 63, 64). The apparently low abundance of proteasome tyrosine phosphorylation is in fact consistent with the low level of total pTyr in cells, which is tightly controlled by the activation status of TKs, spatiotemporal segregation of the phosphorylated proteins and binding partners, as well as active dephosphorylation by PTPs (21, 65). All these mechanisms, as we found in this study, co-operate in regulating Rpt2-Y439 phosphorylation in cells, representing an important novel aspect of proteasome regulation.

Although previous studies using HbYX deletion mutants suggested a non-essential or auxiliary role of Rpt2 tail in 26S proteasome assembly (11, 66), Y439 phosphorylation of wild-type Rpt2 represents a completely different scenario where considerable changes of size and charge are introduced to the tyrosine side chain, which is bound to interfere with RP-CP association as demonstrated in our study. The fact that Y439 phosphorylation does occur on endogenous Rpt2 and is frequently detected by mass spectrometry indicates the existence of a pool of Rpt2 (e.g. in free 19S RP or subcomplexes) outside the fully assembled 26S proteasomes, whose Y439 is accessible to kinases such as Src. This is consistent with the dynamic changes of proteasome composition and the presence of multiple proteasome types in cells (4, 5, 67). In addition, proteasome assembly is also regulated by phosphorylation and cytokine signaling (68–71). It is conceivable that Rpt2-Y439 phosphorylation may participate in these processes as an assembly checkpoint or prevent re-association of Rpt2/RP with CP after they dissociate. Although Rpt2-Y439 is conserved through evolution, its phosphorylation was presumably introduced in metazoans given the apparent lack of canonical tyrosine kinases in yeast (72). Interestingly, the tyrosine residues of the C-terminal HbYX motifs of Rpt3 (Y417) and another proteasome-interacting protein PI31 (Y270) (73) have also been shown to be phosphorylated in human cells (www.phosphosite.org). By the same token, these phosphorylation events may also influence the interaction of Rpt3 and PI31 with the 20S CP, although the regulation and function of these pTyr sites remain to be determined.

The dependence of Rpt2-Y439 phosphorylation on Rpt2 membrane localization was unexpected and also intriguing. The proteasome has been found at various membranes of the cell, and yet how this soluble complex becomes membrane-tethered is unclear (43, 44, 74). In yeast, N-myristoylation of Rpt2 has been regarded as a membrane-targeting signal for localizing the 26S proteasome to the nuclear envelope (38, 39). In human cells, Rpt2 is one of the most myristoylated proteins (31), representing an ideal and perhaps prevalent mechanism for targeting proteasomes to the membrane. Our ongoing research has established that cells expressing the Rpt2-G2A mutant lose most of the membrane-associated proteasomes and that homozygous G2A mutation is embryonic lethal mice. The latter finding illustrates the absolute requirement for localized protein turnover at membranes during embryonic development, although the most relevant functions (substrates) of membrane proteasomes have yet to be confirmed. On the other hand, as a result of Rpt2 N-myristoylation, its C-terminal Y439 can gain access to membrane-localized TKs such as Src. Extracellular signals can then impinge upon membrane-associated proteasomes via RTK/Src and cause changes of the local proteome as suggested by our mass spectrometry study (Fig. 6F). The observation of endogenous Rpt2-Y439 phosphorylation in the developing brain (Fig. 1D) also makes one wonder whether this contributes to neuronal cell migration, synapse formation, etc. that involve drastic reorganization/mobilization of cellular membrane and intracellular organelles. To our knowledge, Rpt2-Y439 phosphorylation is the first evidence of phosphoregulation of membrane-associated proteasomes, which in a broader sense lends weight to the concept of compartmentalized proteasomal degradation.

Membrane localization of Rpt2 is necessary but not sufficient for Y439 phosphorylation, which also relies on Src activity as shown in 293T and H1975 cells. A previous study reported that Src indirectly downregulates proteasome activity by activating the p38 MAPK (75). Our work uncovered that Src directly phosphorylates Rpt2-Y439 and inhibits membrane proteasome activity (Fig. 3D and F) while saracatinib blocks Rpt2-Y439 phosphorylation and promotes proteasomal degradation of membrane-related substrates (Fig. 3E, F). This appears to be a biologically relevant arm of saracatinib-elicited cellular responses, as the Rpt2-Y439F mutation in H1975 cells caused partial resistance to saracatinib and markedly different changes at the proteome level (Figs. 5 and 6). Almost every one of the Group I/II proteins (Fig. 5E) has been found to be ubiquitinated (PhosphoSitePlus). Some of them (as represented by RRas and PAK1) may be substrates of membrane-anchored proteasomes and become more unstable upon saracatinib treatment. For the other proteins such as Legumain as well as those in Groups III and IV, their changes in response to saracatinib were likely to be a secondary effect or independent of membrane proteasome activation. More work is needed to verify which of those saracatinib-downregulated proteins in H1975-WT cells are physiological substrates of membrane-anchored proteasomes and responsible for tumor suppression.

We have also noticed that, in the absence of saracatinib, blocking Y439 phosphorylation did not cause obvious growth defects of the H1975-Y439F cells in vitro or in vivo. This was likely because the Y→F substitution could not fully mimic unphopshorylated wild-type Rpt2 but in effect caused a slight decrease of proteasome activity (e.g. Fig. 1G). Thus, membrane proteasome activity was dampened in both H1975-WT and YF cells but via different mechanisms (Y439 phosphorylation vs. Y439F mutation). However, saracatinib could relieve Src-imposed proteasome inhibition in H1975-WT cells but not in YF cells, hence unmasking the difference between the two. This led to the identification of the 191 proteins differentially regulated by saracatinib between H1975-WT and YF cells (Fig. 5E), though, for the reason discussed above, cautions should be taken when interpreting these data. It is also quite possible that other Src family members or certain RTKs themselves may modify Rpt2-Y439 in different cell types or under different conditions. It would be interesting to examine the impact of Y439 phosphorylation on the therapeutic effects of other TKIs.

In this work, we have confirmed PTPN2 as the first PTP known to bind and regulate the proteasome. Our data suggest that PTPN2 positively regulates proteasome activity through dephosphorylating Rpt2-Y439, and the membrane-associated PTPN2 isoform, TC48, fits particularly well in this scheme. We do not exclude the possibility that TC48 and the other PTPN2 isoforms may regulate different pTyr sites of the proteasome. We also believe that multiple PTPs act in concert to maintain the low level of proteasome tyrosine phosphorylation in the cell, which may be important for cellular fitness. Even for Rpt2-Y439, its phosphorylation still decreased in PTPN2 knockout cells upon Src inhibition (Fig. 4C), albeit more slowly than in control cells. This result indicates the existence of other pY439 phosphatase(s) yet to be identified.

PTPs have emerged as novel drug targets for treating multiple diseases including diabetes, autoimmune disorders and cancer (29, 30, 76). Although PTPN2 inactivates EGFR and Src (54, 77) and has been considered a tumor suppressor (28), recent studies have demonstrated that deletion of PTPN2 in cancer cells or in T cells greatly boosts immunotherapy by upregulating the JAK/STAT pathway (78–80). In addition, variants of the *PTPN2* gene that decrease PTPN2 expression were found to be associated with reduced risk of lung cancer (81). Our data imply that upregulated Rpt2-Y439 phosphorylation following PTPN2 inactivation may exaggerate the impact of activated RTKs/Src on membrane-associated proteasomes and render cancer cells more sensitive to TKIs such as saracatinib. The clinical importance of proteasome tyrosine phosphorylation in cancer treatment warrants further investigation.

## Materials and Methods

### Cell culture, transfection and infection

293T, NCI-H1975, NCI-H3255 and HeLa cells were obtained from American Type Culture Collection (ATCC) and maintained in DMEM (293T, HeLa) or RPMI-1640 (H1975, H3255) with 10% fetal bovine serum (FBS) and Penicillin/Streptomycin (Thermo Fisher). Cells were treated periodically with ciprofloxacin (Sigma) to prevent mycoplasma growth. Transient transfection of cells was performed with the X-tremeGENE 9 reagent (Roche) or polyethylenimine (PEI, Polysciences). Stable cell lines were generated by transduction with retro/lentiviruses followed by puromycin (Thermo Fisher) selection as described (82).

### Plasmids, shRNAs and sgRNAs

All cDNAs of human proteasome subunits with N- or C-terminal Flag tag (except Rpn11) were kindly provided by Dr. Shigeo Murata (University of Tokyo). Rpn11-TBHA was modified from Rpn11-HTBH (41) in the retroviral vector pQCXIP (Clontech). The TBHA tag consists of a biotinylation sequence (“B”), a HA tag and a recognition sequence of the tobacco etch virus protease (TEV, “T”). Rpn1-TBHA, α3-TBHA and Src-TBHA were then generated by replacing Rpn11 with each of the cDNAs. Wild-type Rpt2-IF was made by joining aa. 1-84, Flag tag sequence (DYKDDDDK) and aa. 92-440 of Rpt2 using Gibson Assembly (New England BioLabs), upon which all Rpt2 mutants were built. Myr^Src^-Rpt2-IF-ΔG contains the first 14 aa. of human Src (offered by Dr. Bin Zhao, Life Sciences Institute - Zhejiang University). Myr^Rpt2^-GFPodc was constructed by adding the coding sequence of Rpt2 aa. 1-24 to the front of GFPodc (50) via PCR cloning. Tyrosine kinase cDNAs were from the human kinome cDNA library generously shared by the late Dr. Susan Lindquist (Massachusetts Institute of Technology). PTPN2 (TC48 and TC45) cDNAs were provided by Dr. Xin-Hua Feng (LSI-ZJU). All point mutations were introduced using the QuikChange™ method (Agilent) or Gibson Assembly and verified by sequencing. PTPN2-targeting shRNAs and the control sequence were expressed from the pLKO.1 vector. For simultaneous Rpt2 knockdown and re-expression, pLL3.7 vectors (from Dr. Tyler Jacks, Massachusetts Institute of Technology) were built using a similar strategy described earlier (82). For PTPN2 knockout, sgRNA sequences were inserted into the PX458 or PX459 vectors (Addgene) for transfection into 293T or HeLa cells (69). Sequences of PCR primers and short oligos are provided in Table S2.

### Antibodies and reagents

Anti-Rpt2-pY439 phospho-specific antibodies were raised by immunizing rabbits with the following phospho-peptide: Cys-ENVLYKKQEGTPEGL[pY]L. Positively enriched antibodies were further purified by negative adsorption against immobilized His-SUMO-Rpt2 expressed from *E. coli*. Antibodies generated against the non-phosphorylated version of the above peptide were also affinity-purified and used for anti-Rpt2 immunoprecipitation. Other commercial antibodies are listed in Table S1. The fluorogenic peptide substrate Suc-LLVY-AMC was purchased from UBPBio. MG-132 and all tyrosine kinase inhibitors were purchased from SelleckChem. ATP, cycloheximide and protease inhibitors were bought from Sigma. Fresh working solution of pervanadate was prepared before each experiment by mixing activated sodium orthovanadate (Sigma) with H_2_O_2_ in the dark. The PTPN2 inhibitor Compound 8 (55) was synthesized and generously provided by Dr. Zhong-Yin Zhang (Purdue University).

### Immunoprecipitation, affinity pulldown and immunoblotting

Cells were generally lysed with TBS buffer (50 mM Tris, pH 7.5, 125 mM NaCl, 5-10 mM MgCl_2_) or proteasome lysis/assay buffer (60 mM HEPES, pH 7.6, 50 mM NaCl, 50 mM KCl, 5 mM MgCl_2_, 0.5 mM EDTA, 10% glycerol) supplemented with 0.5% Nonidet P-40, 1 mM ATP, protease inhibitors (1 mM Pefabloc, 1 mM benzamidine hydrochloride, 1 µM Leupetin, 1 µM E-64 and 1 mM phenylmethanesulfonyl fluoride) and phosphatase inhibitors (10 mM NaF, 20 mM β-glycerolphosphate, and 1-10 mM activated orthovanadate). Protein concentration was determined by Bradford protein assay (Bio-Rad). For immunoprecipitation, 0.5-1.0 mg of cell lysate was incubated with 2-4 µg of antibody for 1 h followed by incubation with 8-10 µl of Protein G agarose (Thermo Fisher) for another 30-45 min. For anti-Flag immunoprecipitation of Rpt2-IF, cells were lysed under denaturing conditions (with 1-2% SDS) for complete membrane extraction. Cell lysates were then diluted with the above TBS buffer, sonicated and spin-cleared before mixing with anti-Flag antibody. For streptavidin pulldown, lysates were allowed to bind with 8-15 µl of High Capacity Streptavidin Agarose (Thermo Fisher) for 30-60 min. Beads were washed with the above buffers for 3-4 times. All incubation and washing were performed at 4°C. Bound proteins were eluted by boiling in Laemmli sample buffer, separated on SDS-PAGE and transferred to nitrocellulose membranes for immunoblotting. Exposed films were scanned and band intensities were quantified by ImageJ (https://imagej.nih.gov/ij/). For native gel analysis, non-denatured cell lysates were separated on NativePAGE™ 3-12% Bis-Tris gel (Thermo Fisher) following manufacturer’s procedure. ATP (1 mM) was supplemented in gel running buffer and electrophoresis was carried out in an ice-water bath. Proteins were transferred to nitrocellulose membranes at 400 mA in a Tris-glycine transfer buffer containing 1% methanol at 4°C for 4 hrs.

### Analysis of proteasome assembly and activity

Proteasome activity assays using the fluorogenic peptide substrates were performed according to the standard protocol (83) with a TECAN infinite M200 Pro multi-well plate reader (TECAN). Protease inhibitors were omitted from cell lysis buffer if cell lysates were directly used for the peptidase assay. To monitor proteasome activity in cells, Myr^Rpt2^-GFPodc was cloned into the pQCXIP vector, with the original CMV promoter replaced by the UbiC promoter. Retroviruses were packaged and used for stable transduction of 293T cells. Then single clones were isolated and those with optimal expression levels of the reporter proteins that were highly sensitive to proteasomal degradation were chosen. GFP level was determined by anti-GFP western blot.

### Fluorescence microscopy

HeLa cells grown on glass coverslips were transfected with Rpt2-IF and GFP-TC48 as indicated. Cells were fixed, permeabilized and blocked as reported (84). Primary and secondary antibodies used are listed in Table S1. Cells were then stained with DAPI (Yeasen, China), and coverslip mounted with Mowiol (Sigma). Z-stack images were taken with a spinning disk confocal microscope (Andor) under a 100X oil lens, and analyzed with the Metamorph software package.

### Membrane/cytosol fractionation

Cell fractionation was adapted from a protocol of the Lamond Lab (http://www.lamondlab.com). 293T cells were scraped off from culture plates into hypotonic, detergent-free lysis buffer (5 mM HEPES, pH 7.9, 5 mM MgCl_2_, 1 mM ATP) supplemented with protease and phosphatase inhibitors. After incubation on ice for 5 min, swollen cells were broken with a Dounce homogenizer with a tight-fitting pestle. A small aliquot was saved as total protein control, and the rest of sample was centrifuged at 2,800 x g, 4°C for 10 min to remove unbroken cells, nuclei and large cell debris. The supernatant was further centrifuged at 21,500 x g, 4°C for 20 min. The new supernatant was saved as cytosolic fraction, while the pellet (mostly membranes) was washed twice with the same hypotonic buffer. The remaining pellet was then dissolved in TBSN buffer (TBS + 1% NP-40) and centrifuged at 21,500 x g, 4°C for 10 min. The soluble proteins extracted were considered the membrane fraction. NP-40 was also added to the cytosolic fraction (1% final), and both cytosolic and membrane fractions were quantified by Bradford protein assay. Equal amounts (10-20 µg) of proteins from each fraction were used for western blot.

### Protein purification and *in vitro* kinase/phosphatase assays

Recombinant His-SUMO-Rpt2 (WT or Y439F) proteins were purified from *E.coli* (BL21-CodonPlus (DE3)-RIPL, Agilent) as previously described (69). The PTP domain of human PTPN2 (aa. 1-314, WT or C216S) with a N-terminal 8x His tag was expressed from the pSJ2 vector (84) in BL21 *E.coli* and purified by Ni-NTA resin (Thermo Fisher). Full-length human Src kinase (WT or K298M) with a C-terminal TBHA tag was stably expressed in 293T cells. Src-TBHA was purified by streptavidin pulldown followed by extensive washes, and Src was cleaved off by TEV in TBS buffer. TEV (with a N-terminal His tag) was removed by Ni-NTA resin, and 10% glycerol was added to purified Src before storage at −80°C.

For *in vitro* kinase assay, recombinant His-SUMO-Rpt2 proteins were incubated with purified Src in kinase buffer (50 mM HEPES, pH 7.4, 5 mM MgCl_2_, 1 mM ATP, 40 µg/ml BSA) at 30°C for 1 hr. For *in vitro* phosphatase assay, Rpt2-IF was isolated by anti-Flag immunoprecipitation from cells pre-treated with pervanadate (0.1 mM, 30 min). After extensive washes with the phosphatase buffer (50 mM Tris, 50 mM Bis-Tris, pH 7.5, 100 mM NaOAc, 2 mM DTT)(55), purified PTPN2 protein was added and incubated with Rpt2-IF (substrate) at 30°C for 1 hr. For both kinase and phosphatase assays, reactions were terminated by boiling the samples in 1X Laemmli sample buffer and pY439 levels were examined by western blot.

### Colony formation assay

H1975-WT and YF cells were counted and seeded in triplicates in 12-well plates (1,000 cells/well). Different concentrations of saracatinib were added the next day. After 48 hrs, cells were gently but extensively washed to remove the drug. Fresh medium was added and cells were allowed to grow for another 7 days under normal culture conditions. The plate was then washed with PBS and fix-stained with 0.5% crystal violet. Each well was thoroughly washed with distilled water, dried and photographed. Crystal violet of the stained cells was then dissolved with ethanol plus 1% (v/v) of concentrated HCl, and Abs_590_ was measured on a Tecan multiwell plate reader.

### Proteomic analysis

H1975-WT and YF cells treated with DMSO or saracatinib (250 nM, 24 hrs) were harvested in SDT buffer (4% SDS, 0.1M DTT, 0.1M Tris, pH 7.6). The filter-aided sample preparation method (FASP)(85) was used to prepare digested samples for protein identification with 200 µg of each sample. After reduction and alkylation, the solution was buffer-exchanged with 50 mM NH_4_HCO_3_ (pH 8.0–8.5) using 30 kDa molecular weight cut-off Amicon Spin Tube (Millipore, MA). For digestion, 1 µg of sequencing-grade modified trypsin (Promega, USA) was added to each solution, followed by a 37°C overnight incubation. Prior to mass spec analysis, the peptide preparations were desalted using Zip-tip C18 cartridges (Millipore, MA) and reconstituted with 0.1% formic acid (FA).

Mass spectrometry experiments were performed using a Q Exactive HF-X instrument (Thermo Fisher) coupled with an UltiMate 3000 nano LC. Mobile phase A and B were water and 80% acetonitrile, respectively, with 0.1% formic acid. Protein digests were loaded directly onto a C18 PepMap EASYspray column (part number ES803; Thermo Fisher Scientific) at a flow rate of 300 nl/min. All samples were separated using a linear gradient of 2–40% B over 38 min. Survey scans of peptide precursors were performed from 350 to 1500 m/z at 60,000 FWHM resolution with a 1 × 10^6^ ion count target and a maximum injection time of 20 ms. The instrument was set to run in top-speed mode with 1-s cycles for the survey and the MS/MS scans. After a survey scan, tandem MS was then performed on the most abundant precursors exhibiting a charge state from 2 to 7 of greater than 5 × 10^4^ intensity by isolating them in the quadrupole at 1.6 Da. Higher energy collisional dissociation (HCD) fragmentation was applied with 30% collision energy and resulting fragments detected in the Orbitrap detector at a resolution of 15,000. The maximum injection time limited was 45 ms and dynamic exclusion was set to 45 s with a 10-ppm mass tolerance around the precursor.

The raw data acquired were imported into the Maxquant software 1.6.0.1 (Max Planck Institute of Biochemistry, Germany) to identify and quantify the proteins (86). The following parameters were used: trypsin for enzyme digestion; oxidation of methionine as variable modification; main search peptide of 10 ppm; a fragment mass tolerance of 0.02 Da. The human reference database was downloaded from UniprotKB and used as the searching database. The identification criterion for peptides was a <1% FDR value according to the database searching. Label-free quantification was enabled. Raw data of mass spectrometry can be accessed via the following link (http://uaalink.com/download/Guo_FASP.zip).

### Functional enrichment and protein-protein interaction analysis

PANTHER was used for Gene Ontology (GO) enrichment analysis (http://www.pantherdb.org/; version 15.0) (87). Statistical significance was assessed by the Fisher’s exact test. Multi-comparisons were adjusted by the Benjamini and Hochberg method (88) and alpha was set to 0.05, with all genes in *Homo sapiens* reference genome set as the background (the statistical overrepresentation method provided by PANTHER). The visualizations for GO results were done in Rstudio (Version 4.0.0), with ggplot2 (Version 3.3.1), ggsci (Version 2.9), ggpubr (Version 0.3.0), and RColorBrewer (Version 1.1-2) packages.

The protein-protein interaction (PPI) network of the differentially regulated proteins was analyzed by STRING (https://string-db.org; version 11.0) (89), with interaction confidence set to 0.40 and MCL clustering (MCL inflation parameter set to 3).

### Animal studies

All animal studies were conducted in full compliance with policies of the Institutional Animal Care and Use Committee (IACUC) of Zhejiang University. Six-week old female nu/nu mice were purchased from SLAC Laboratory Animal Co., Ltd (Shanghai, China) and housed at the Laboratory Animal Center at Zhejiang University. Each mouse was injected subcutaneously at the flank with 2.0 x 10^6^/100 µl of H1975 cells, which were resuspended in PBS and mixed 1:1 (v/v) with Matrigel (Corning, #354234). Two weeks later, tumor-bearing mice were randomized and separated to receive DMSO or saracatinib (25 mg/kg/d) treatment. Saracatinib was dissolved in DMSO and diluted in PEG300 as recommended by the vendor (Selleckchem), and administered daily by oral gavaging. Tumor size was measured every other day with a digital caliber, and tumor volume was calculated as V (mm^3^) = ½ (width^2^ x length). At the end of the study, mice were euthanized and tumors were resected out, weighed and immediately frozen in liquid nitrogen then stored at −80°C. Tumor tissues were later Dounce-homogenized in hypotonic lysis buffer (containing protease and phosphatase inhibitors) and sonicated. NP-40 (1% final) was then added to facilitate membrane protein extraction. Tissue lysates were cleared by centrifugation and quantified by BCA protein assay (Thermo Fisher) for western blot analysis.

Pregnant wild-type Sprague Dawley rats were also purchased from SLAC Laboratory Animal Co., Ltd (Shanghai, China). E15.5 embryos were dissected, and forebrain/midbrain tissues were homogenized in TBS buffer supplemented with 0.1% SDS, protease/phosphatase inhibitors, ATP and MgCl_2_. Tissue homogenates were further sonicated before anti-Rpt2 immunoprecipitation.

### General methods for data analysis

Structural models were generated with PyMol. Sequence alignment was performed by ClustalW. Gel analysis was done by ImageJ. Quantitative results are shown as mean ± S.D. and statistical analyses were done with GraphPad Prism.

## Supporting information

Table S3 overlapping proteins

Table S4 D-scores

Table S5 Downregulated proteins

Table S6 Upregulated proteins

Table S7 Diff. reg WT vs. YF

## Acknowledgements

We thank Drs. Xin-Hua Feng, Hai Song, Bin Zhao (Life Sciences Institute, Zhejiang University, LSI-ZJU), Zhong-Yin Zhang (Purdue University), Shigeo Murata (The University of Tokyo) and Susan Lindquist (Massachusetts Institute of Technology) for crucial constructs, cell lines and reagents. We are grateful for the critical and encouraging comments from Drs. Kun-Liang Guan, Anne-Claude Gingras, Benjamin Neel, Tony Hunter and Tony Tiganis. We appreciate the technical assistance from Xiaorui Jiang, Fei Zhang and the Core Facilities of LSI-ZJU. X.G. was supported by Natural Science Foundation of China (31671391, 31870762), Zhejiang Natural Science Foundation (LR18C050001), Fundamental Research Funds for the Central Universities (2016QN81011), and the startup funding from Zhejiang University. B.Y. was funded by Natural Science Foundation of China (91953103). Z.W. was supported by Natural Science Foundation of China (31671039) and National Key Research and Development Plan of the Ministry of Science and Technology of China (2016YF0501000). L.H. was funded by NIH (R01GM074830).

## Supplementary Figures

**Supplementary Fig. S1.**
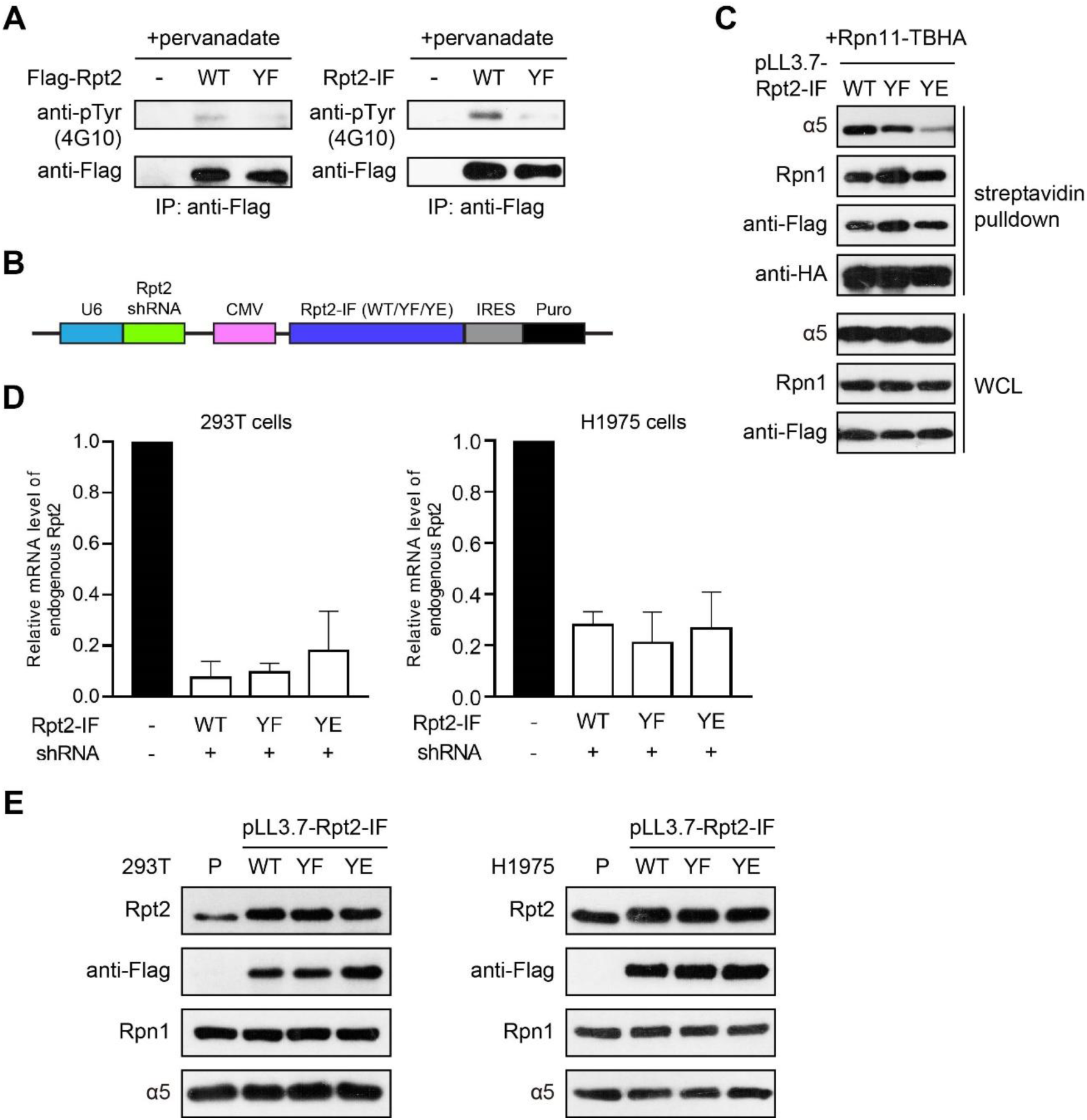
Expression and characterization of Rpt2-Y439 mutants. **A.** N-terminal Flag-tagged (left) and internal Flag-tagged (right) Rpt2-WT/Y439F were expressed in 293T cells. After pervanadate treatment and anti-Flag immunoprecipitation, Rpt2 tyrosine phosphorylation was determined by western blot. **B.** Schematic of the pLL3.7-Rpt2 construct for simultaneously knocking down endogenous Rpt2 and re-expressing Rpt2-IF variants. **C.** 293T cells were stably transduced with pLL3.7-Rpt2-IF-WT/Y439F/Y439E. Rpn11-TBHA was transiently expressed in these cells. Following streptavidin pulldown, co-isolation of the different proteasome subunits was shown by western blot. **D.** qPCR measurement of endogenous Rpt2 mRNA in 293T and H1975 cells stably transduced with pLL3.7-Rpt2 shown in (C), n = 3 for each cell type. **E.** Western blot verification of exogenous Rpt2-IF expression in the above 293T and H1975 cells.

**Supplementary Fig. S2.**
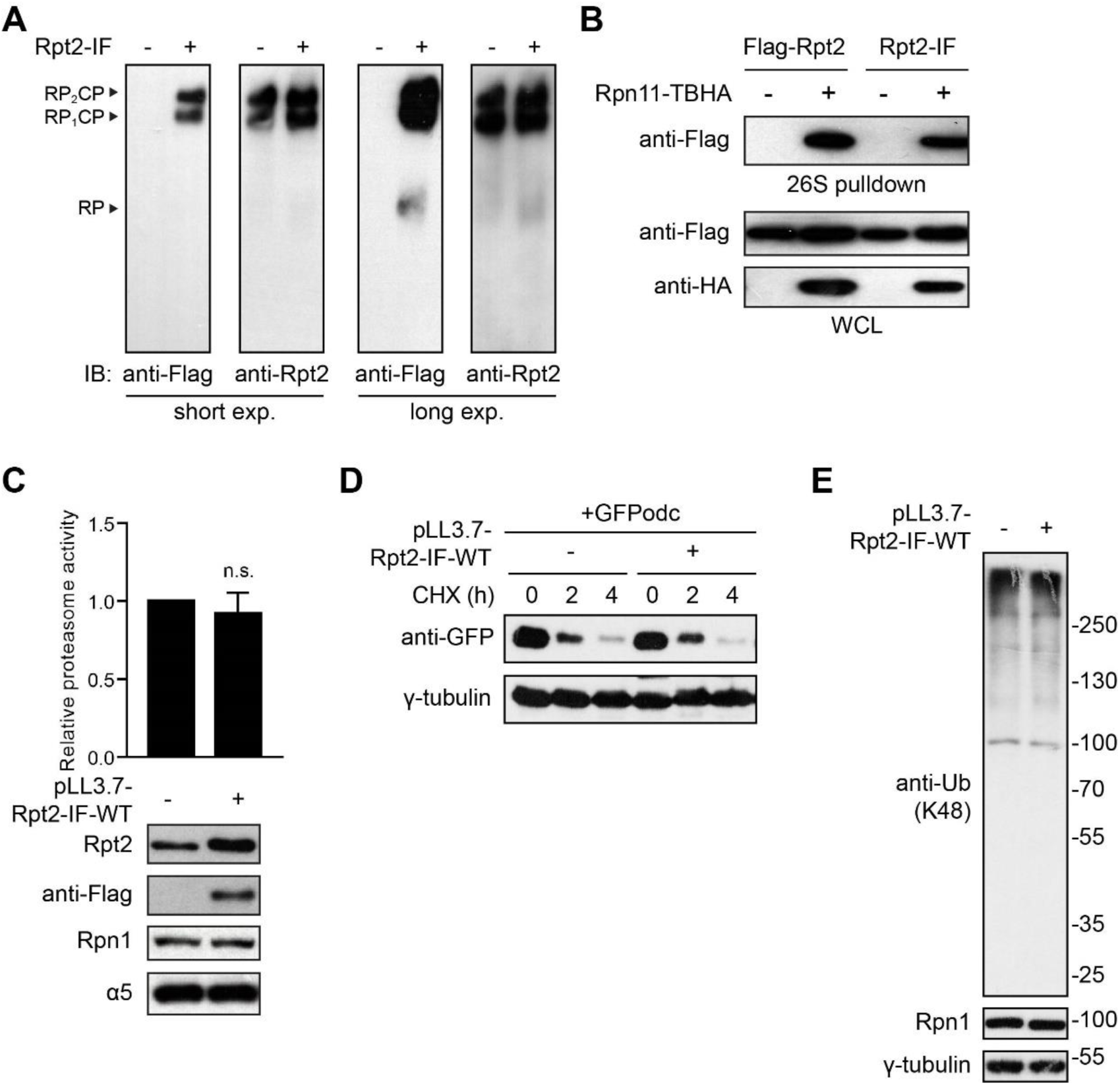
The internal Flag tag does not interfere with Rpt2 function. **A.** 293T cells were transfected with or without Rpt2-IF, and proteasome complexes from whole cell lysates were separated by native gel and visualized by immunoblotting. **B.** Flag-Rpt2 and Rpt2-IF were co-transfected with vector (-) or Rpn11-TBHA into 293T cells. Following streptavidin pulldown, proteasome-associated Rpt2 was shown by western blot. **C.** Parental (-) and pLL3.7-Rpt2-IF-WT transduced (+) 293T cells as in Supplementary Fig. 1D, E were assessed for proteasome activity using Suc-LLVY-AMC as substrate (n = 3). n.s., not significant. **D.** The same 293T cells as in (C) were transfected with GFPodc and treated with CHX as indicated. GFPodc degradation was determined by western blot. **E.** The same 293T cells as in (C) were lysed in the presence of N-Ethylmaleimide (NEM, 10 mM). K48-linked polyubiquitinated proteins from whole cell lysates were probed.

**Supplementary Fig. S3.**
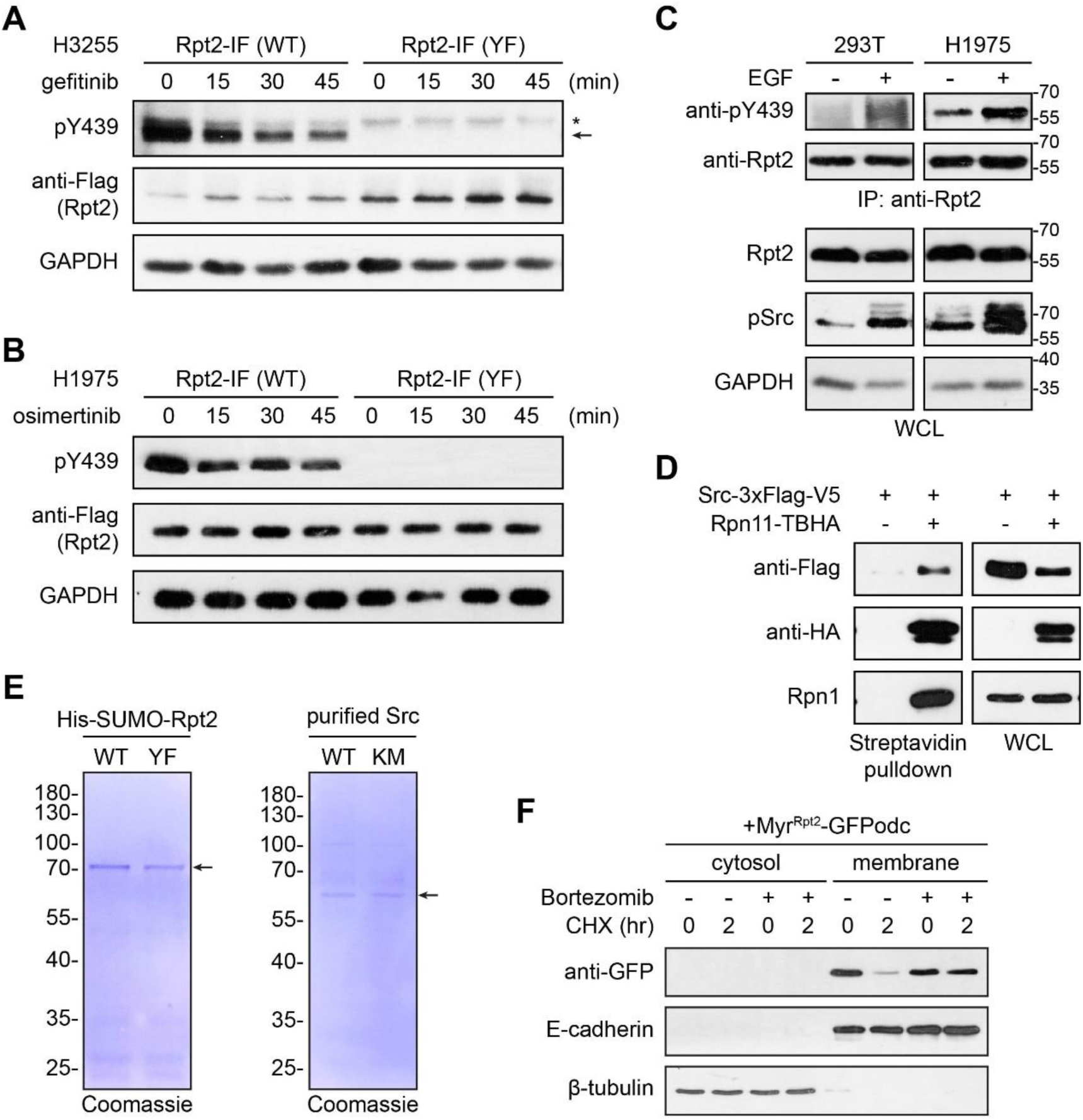
EGFR/Src regulate Rpt2-Y439 phosphorylation. **A and B.** H3255 (A) and H1975 (B) NSCLC cells were stably transduced with pLL3.7-Rpt2-IF-WT or Y439F. Cells were treated with 10 µM gefitinib or osimertinib, respectively, for the indicated time course. Whole cell lysates were probed with the indicated antibodies. In (A), the arrow points to pY439 of Rpt2-IF, while the asterisk indicates a non-specific band. **C.** 293T and H1975 cells were serum-starved for 24 hrs then treated with recombinant hEGF (50 ng/ml) for 10 min. Cells were immediately lysed and the same amount of proteins were subjected to anti-Rpt2 immunoprecipitation. Samples were probed with the indicated antibodies. **D.** Src-3xFlag-V5 was co-transfected with vector or Rpn11-TBHA into 293T cells. Physical binding of Src to the proteasome was determined by western blot after streptavidin pulldown. **E.** Coomassie staining of His-SUMO-Rpt2 proteins (WT or Y439F) purified from *E. coli* (left) and Src kinase purified from 293T cells stably expressing Src-WT/K298M-TBHA (right). **F.** A single clone of 293T cells stably expressing Myr^Rpt2^-GFPodc was treated with cycloheximide (CHX, 50 µg/ml) or bortezomib (1 µM, 4 hrs) or both as indicated. The cytosolic and solubilized membrane fractions were probed with the indicated antibodies.

**Supplementary Fig. S4.**
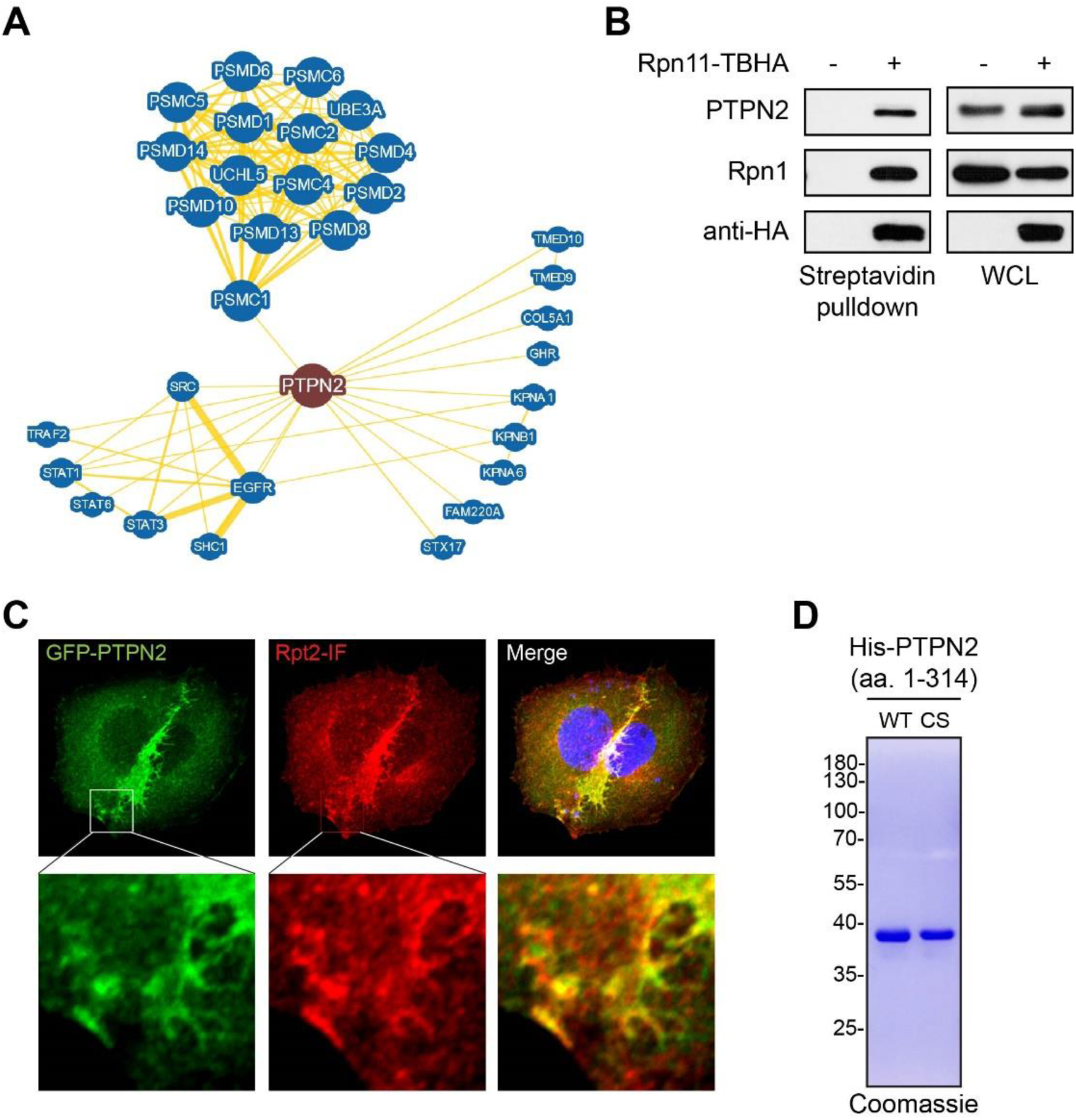
PTPN2 is a proteasome-interacting phosphatase. **A.** PTPN2 interactome from BioGRID. Rpt2/PSMC1 is the only proteasome subunit that has been shown to interact with PTPN2 by more than two studies. **B.** 293T cells stably expressing Rpn11-TBHA or TBHA only (-) were lysed for streptavidin pulldown. Endogenous PTPN2 readily co-purified with the proteasome as shown by western blot. **C.** HeLa cells were co-transfected with GFP-TC48 and Rpt2-IF (WT), immunostained with anti-Flag antibody and imaged with a con-focal microscope. Upper panel: 500 x 500 pixels (1 pixel = 0.1375 um). Zoom-in of the indicated areas are show in the bottom panel (100×100 pixels). **D.** Coomassie staining showing successful purification of the PTPN2 phosphatase domain from *E. coli*. CS, C216S.

**Supplementary Fig. S5.**
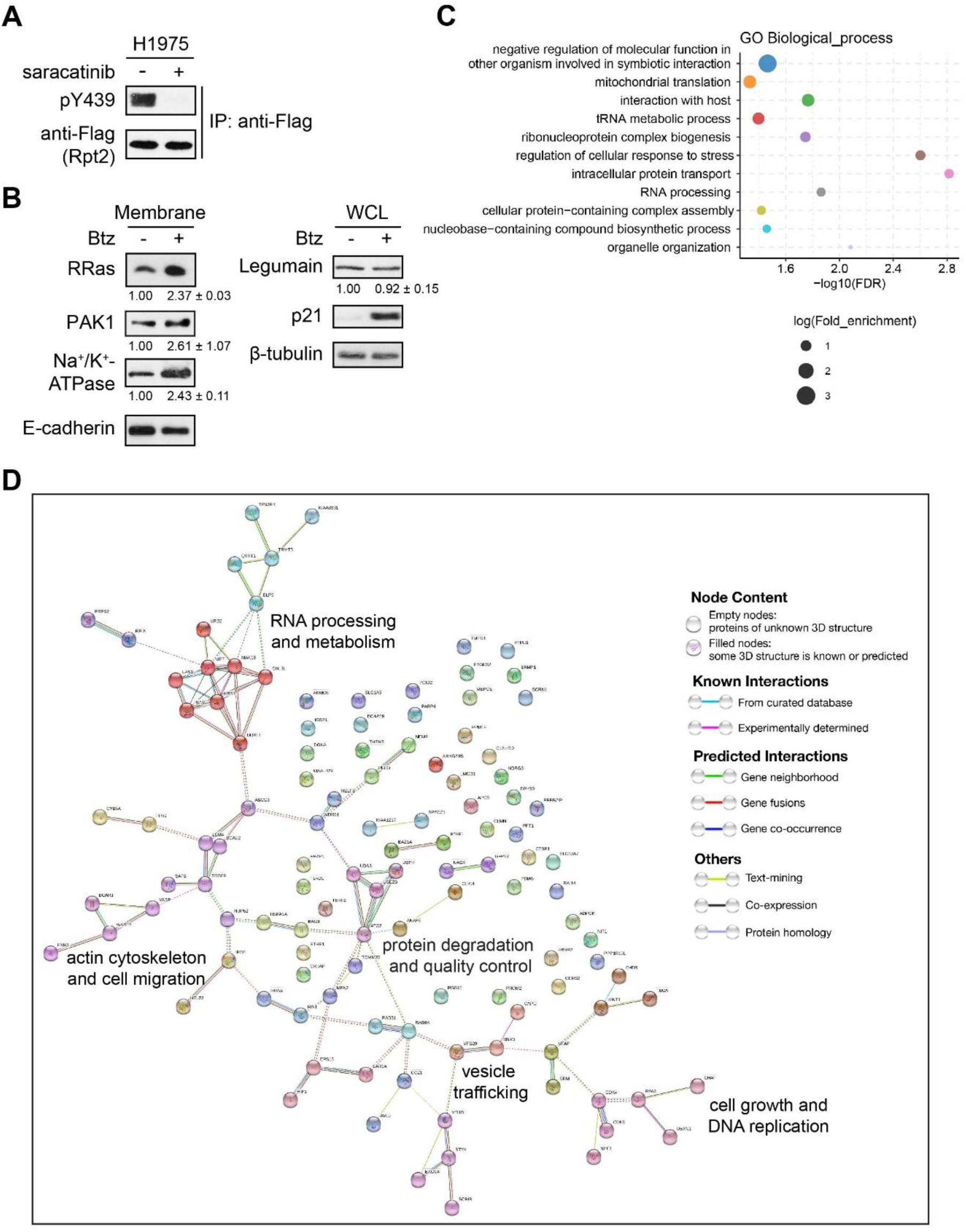
Saracatinib induces distinct proteomic changes in H1975-WT and YF cells. **A.** H1975-WT cells (stably transduced with pLL3.7-Rpt2-IF-WT) were treated with DMSO or saracatinib (10 µM, 1 hr). Cell lysates were probed with the antibodies shown. **B.** H1975 cells were treated with DMSO or bortezomib (1 µM, 4 hrs). Membrane fractions or whole cell lysates were probed with the indicated antibodies. Relative protein levels from 3 independent experiments were quantified and shown below each blot. **C.** GO term (biological process) analysis of the 191 differentially regulated proteins between H1975-WT and YF cells in response to saracatinib treatment. Only first-level terms in each functional hierarchy (indicated by a different color) with FDR < 0.05 are shown. **D.** The protein-protein interaction network of the 191 differentially regulated proteins. Color codes are defined by MCL clustering (clustering parameter set to 3).

**Table S1.**
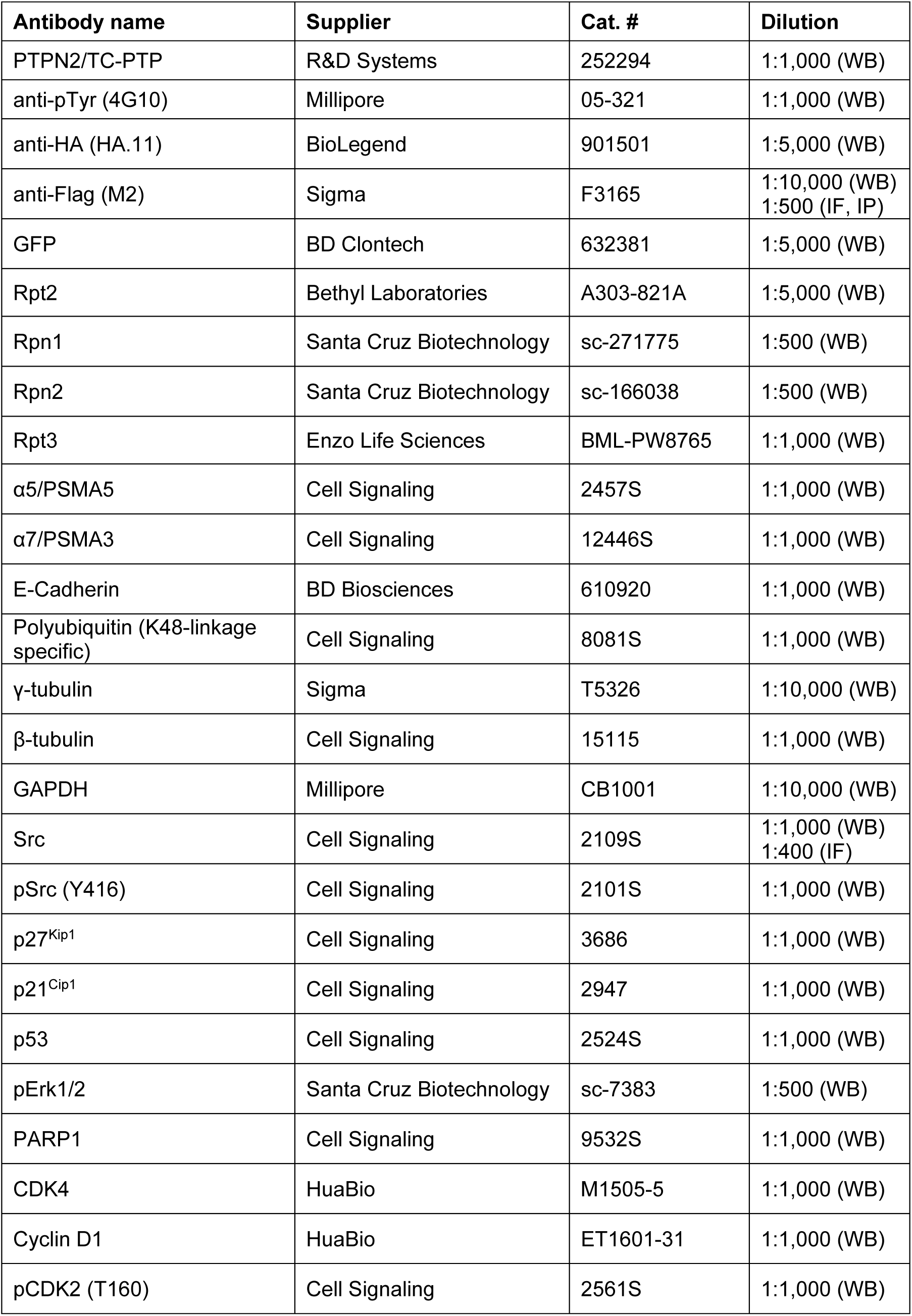
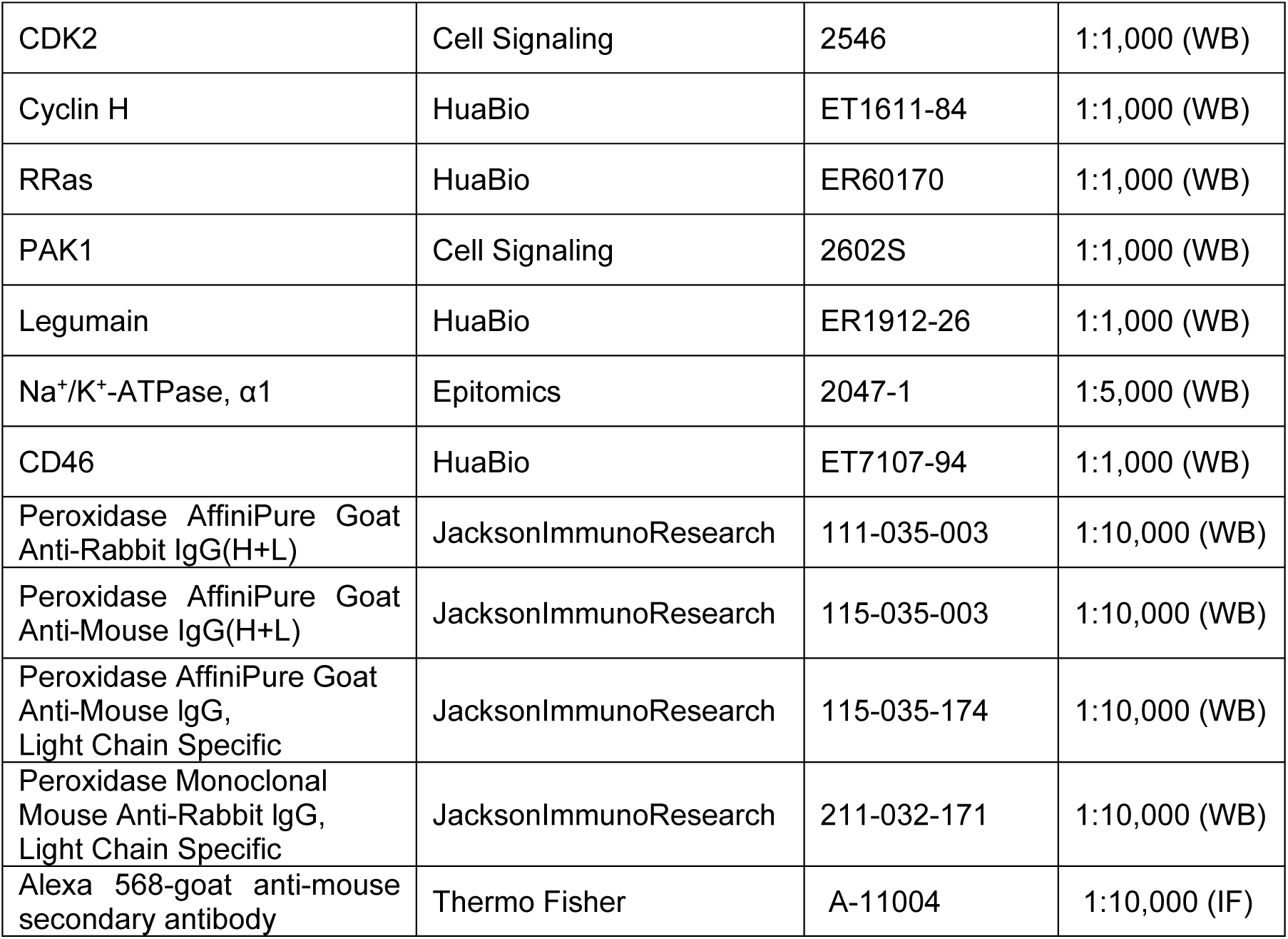
Commercial antibodies used in this study.

| **Antibody name** | **Supplier** | **Cat. #** | **Dilution** |
| --- | --- | --- | --- |
| PTPN2/TC-PTP | R&D Systems | 252294 | 1:1,000 (WB) |
| anti-pTyr (4G10) | Millipore | 05-321 | 1:1,000 (WB) |
| anti-HA (HA.11) | BioLegend | 901501 | 1:5,000 (WB) |
| anti-Flag (M2) | Sigma | F3165 | 1:10,000 (WB)<br>1:500 (IF, IP) |
| GFP | BD Clontech | 632381 | 1:5,000 (WB) |
| Rpt2 | Bethyl Laboratories | A303-821A | 1:5,000 (WB) |
| Rpn1 | Santa Cruz Biotechnology | sc-271775 | 1:500 (WB) |
| Rpn2 | Santa Cruz Biotechnology | sc-166038 | 1:500 (WB) |
| Rpt3 | Enzo Life Sciences | BML-PW8765 | 1:1,000 (WB) |
| α5/PSMA5 | Cell Signaling | 2457S | 1:1,000 (WB) |
| α7/PSMA3 | Cell Signaling | 12446S | 1:1,000 (WB) |
| E-Cadherin | BD Biosciences | 610920 | 1:1,000 (WB) |
| Polyubiquitin (K48-linkage specific) | Cell Signaling | 8081S | 1:1,000 (WB) |
| γ-tubulin | Sigma | T5326 | 1:10,000 (WB) |
| β-tubulin | Cell Signaling | 15115 | 1:1,000 (WB) |
| GAPDH | Millipore | CB1001 | 1:10,000 (WB) |
| Src | Cell Signaling | 2109S | 1:1,000 (WB)<br>1:400 (IF) |
| pSrc (Y416) | Cell Signaling | 2101S | 1:1,000 (WB) |
| p27<sup>Kip1</sup> | Cell Signaling | 3686 | 1:1,000 (WB) |
| p21<sup>Cip1</sup> | Cell Signaling | 2947 | 1:1,000 (WB) |
| p53 | Cell Signaling | 2524S | 1:1,000 (WB) |
| pErk1/2 | Santa Cruz Biotechnology | sc-7383 | 1:500 (WB) |
| PARP1 | Cell Signaling | 9532S | 1:1,000 (WB) |
| CDK4 | HuaBio | M1505-5 | 1:1,000 (WB) |
| Cyclin D1 | HuaBio | ET1601-31 | 1:1,000 (WB) |
| pCDK2 (T160) | Cell Signaling | 2561S | 1:1,000 (WB) |
| CDK2 | Cell Signaling | 2546 | 1:1,000 (WB) |
| Cyclin H | HuaBio | ET1611-84 | 1:1,000 (WB) |
| RRas | HuaBio | ER60170 | 1:1,000 (WB) |
| PAK1 | Cell Signaling | 2602S | 1:1,000 (WB) |
| Legumain | HuaBio | ER1912-26 | 1:1,000 (WB) |
| Na<sup>+</sup>/K<sup>+</sup>-ATPase, α1 | Epitomics | 2047-1 | 1:5,000 (WB) |
| CD46 | HuaBio | ET7107-94 | 1:1,000 (WB) |
| Peroxidase AffiniPure Goat Anti-Rabbit IgG(H+L) | JacksonImmunoResearch | 111-035-003 | 1:10,000 (WB) |
| Peroxidase AffiniPure Goat Anti-Mouse IgG(H+L) | JacksonImmunoResearch | 115-035-003 | 1:10,000 (WB) |
| Peroxidase AffiniPure Goat Anti-Mouse IgG, Light Chain Specific | JacksonImmunoResearch | 115-035-174 | 1:10,000 (WB) |
| Peroxidase Monoclonal Mouse Anti-Rabbit IgG, Light Chain Specific | JacksonImmunoResearch | 211-032-171 | 1:10,000 (WB) |
| Alexa 568-goat anti-mouse secondary antibody | Thermo Fisher | A-11004 | 1:10,000 (IF) |

**Table S2.**
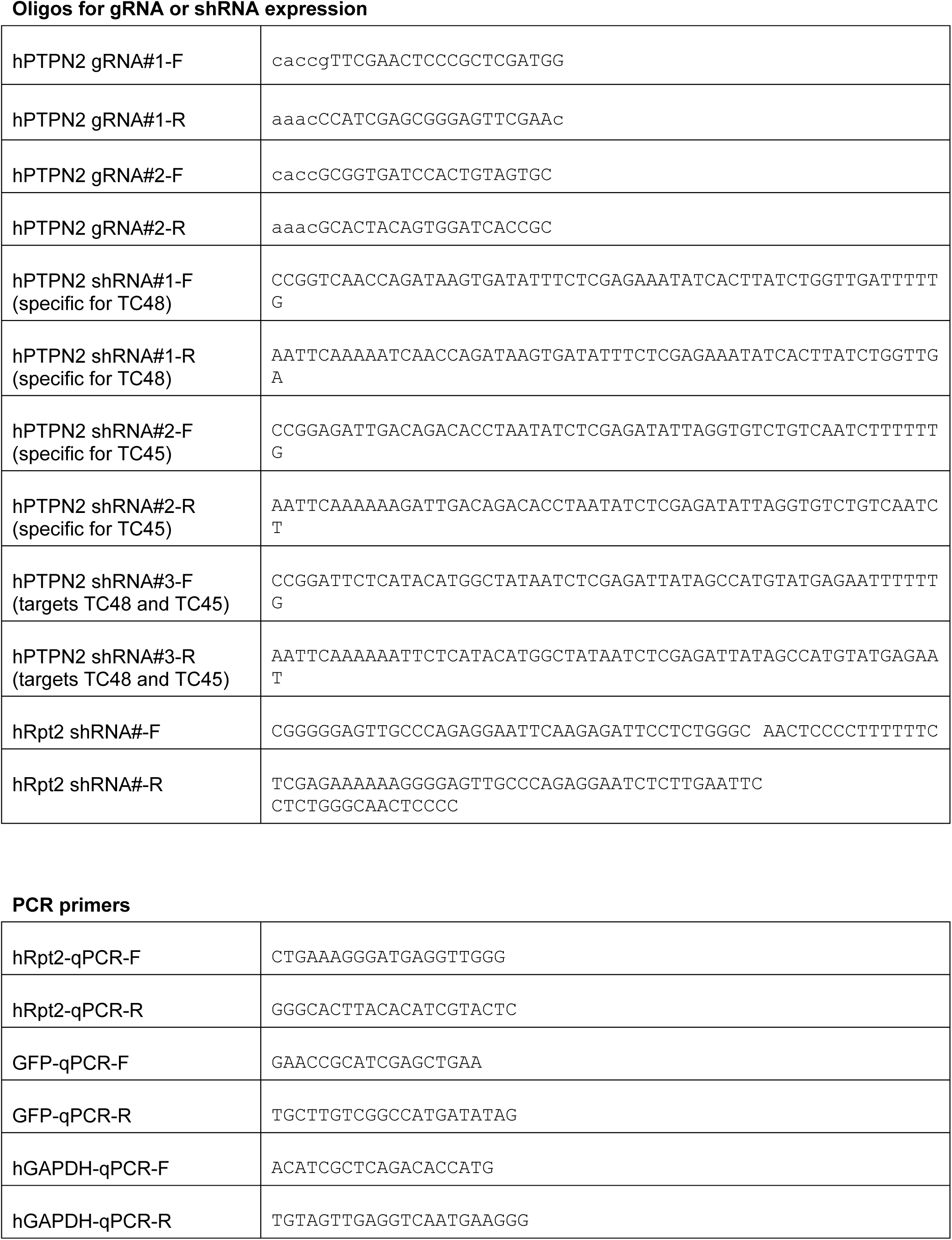
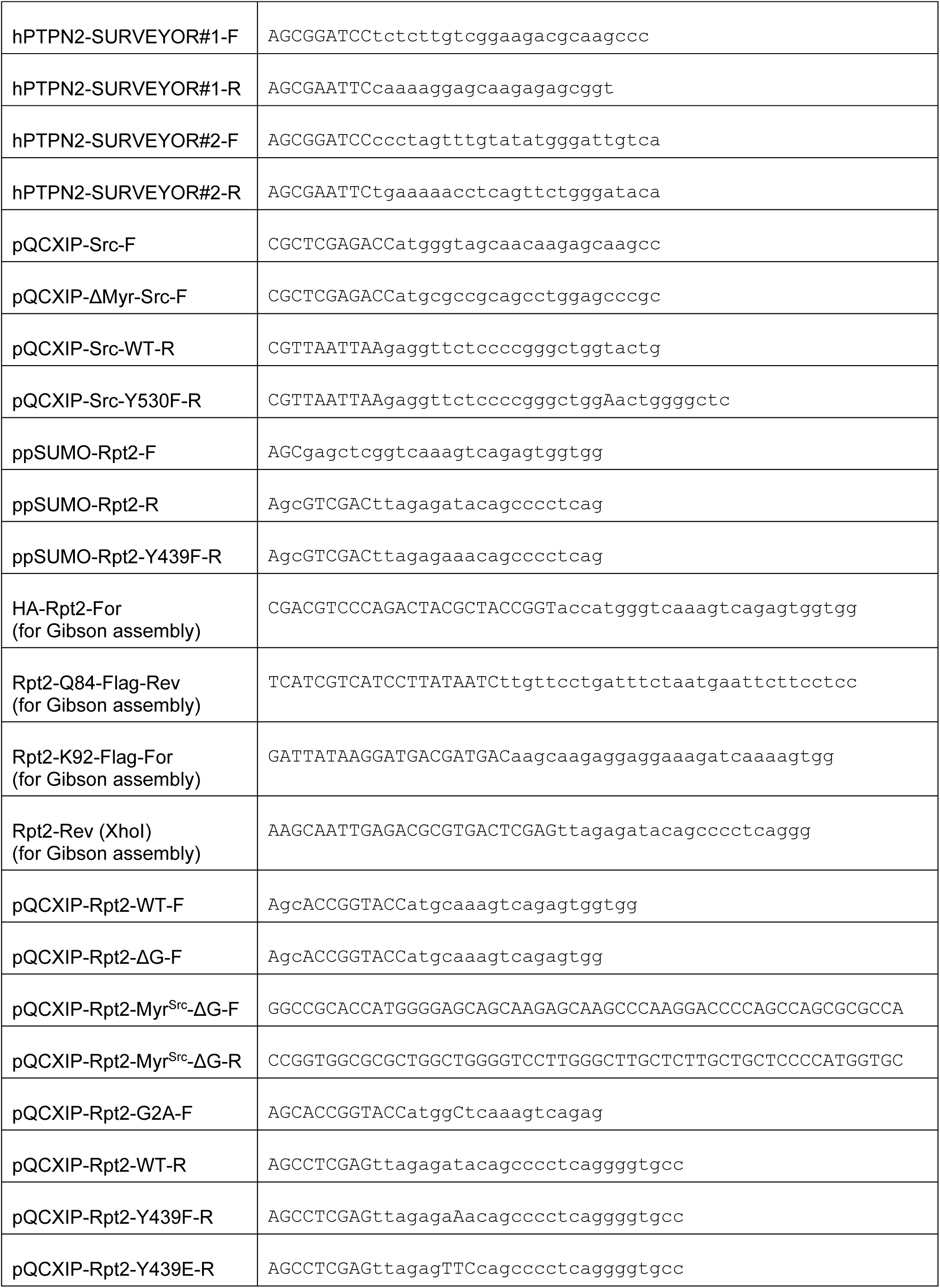
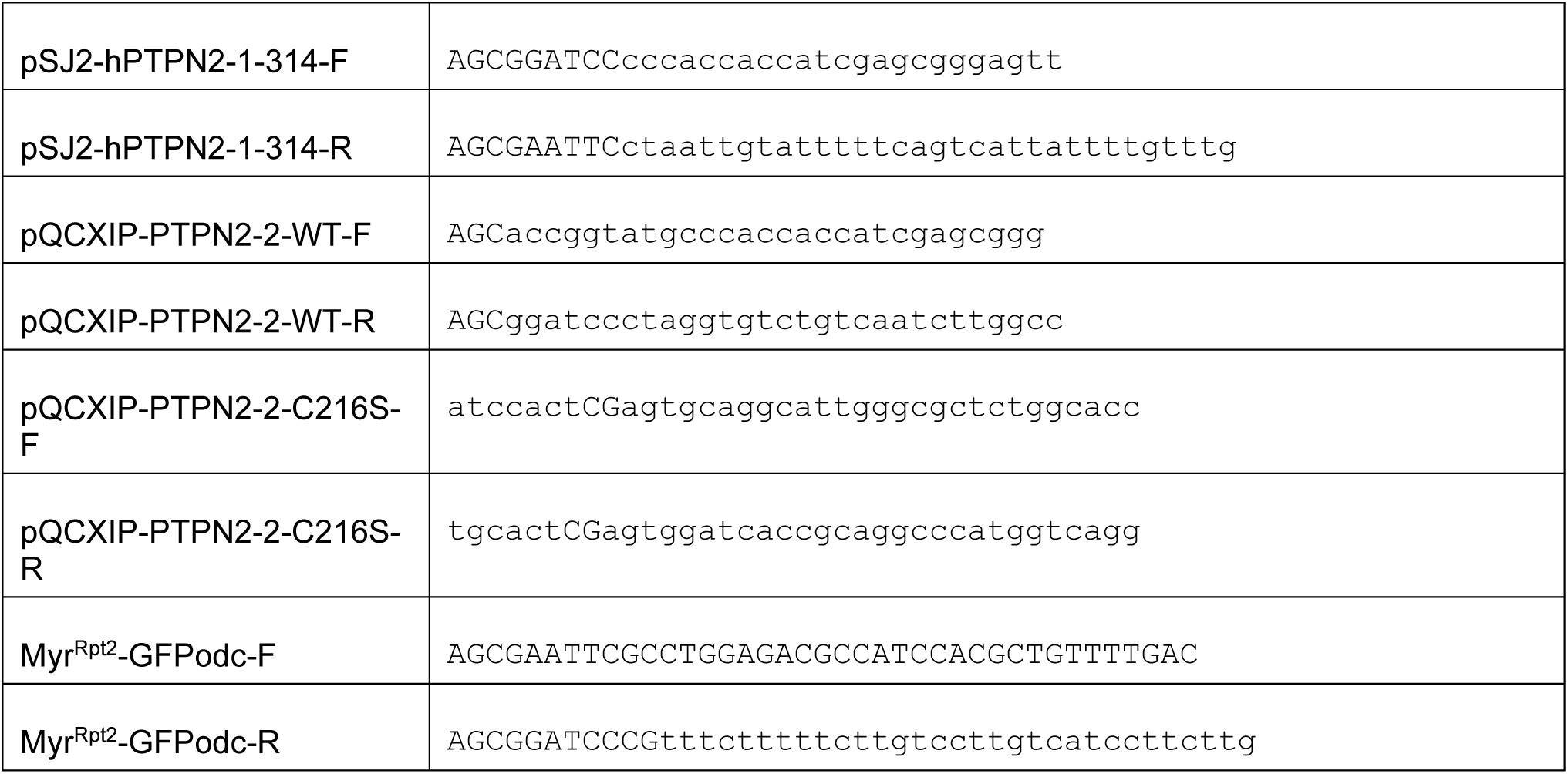
Primer and oligonucleotide sequences.

**Table S3.**

List of all overlapping proteins identified by mass spectrometry in 3 biological repeats. Data from repeat #1, 2, 3 are labeled with yellow, green and blue, respectively, in the header row.

**Table S4.**

D scores of quantifiable proteins across all 3 biological repeats.

**Table S5.**

Proteins downregulated by saracatinib in at least 2 of 3 biological repeats. Proteins annotated as membrane-localized (including plasma membrane, endosome, lysosome, Golgi apparatus, endoplasmic reticulum, peroxisome, nuclear envelope, mitochondria and secreted/exsosome) are indicated.

**Table S6.**

Proteins upregulated by saracatinib in at least 2 of 3 biological repeats.

**Table S7.**

Proteins that were differentially regulated by saracatinib between H1975-WT and YF cells.

## Notes

http://uaalink.com/download/Guo_FASP.zip

### Competing Interest Statement

The authors have declared no competing interest.

## References

1. Finley D & Prado MA (2019) The Proteasome and Its Network: Engineering for Adaptability. Cold Spring Harbor perspectives in biology.

2. Schmidt M & Finley D (2014) Regulation of proteasome activity in health and disease. Biochimica et biophysica acta 1843(1):13–25.

3. Collins GA & Goldberg AL (2017) The Logic of the 26S Proteasome. Cell 169(5):792–806.

4. Murata S, Yashiroda H, & Tanaka K (2009) Molecular mechanisms of proteasome assembly. Nat Rev Mol Cell Biol 10(2):104–115.

5. Budenholzer L, Cheng CL, Li Y, & Hochstrasser M (2017) Proteasome Structure and Assembly. Journal of molecular biology 429(22):3500–3524.

6. Finley D (2009) Recognition and processing of ubiquitin-protein conjugates by the proteasome. Annu Rev Biochem 78:477–513.

7. Bard JAM, et al. (2018) Structure and Function of the 26S Proteasome. Annu Rev Biochem 87:697–724.

8. Eisele MR, et al. (2018) Expanded Coverage of the 26S Proteasome Conformational Landscape Reveals Mechanisms of Peptidase Gating. Cell Rep 24(5):1301–1315.e1305.

9. Smith DM, et al. (2007) Docking of the proteasomal ATPases’ carboxyl termini in the 20S proteasome’s alpha ring opens the gate for substrate entry. Mol Cell 27(5):731–744.

10. Park S, et al. (2013) Reconfiguration of the proteasome during chaperone-mediated assembly. Nature 497(7450):512–516.

11. Park S, et al. (2009) Hexameric assembly of the proteasomal ATPases is templated through their C termini. Nature 459(7248):866–870.

12. Kumar B, Kim YC, & DeMartino GN (2010) The C terminus of Rpt3, an ATPase subunit of PA700 (19 S) regulatory complex, is essential for 26 S proteasome assembly but not for activation. J Biol Chem 285(50):39523–39535.

13. Livneh I, Cohen-Kaplan V, Cohen-Rosenzweig C, Avni N, & Ciechanover A (2016) The life cycle of the 26S proteasome: from birth, through regulation and function, and onto its death. Cell Res 26(8):869–885.

14. Hoeller D & Dikic I (2009) Targeting the ubiquitin system in cancer therapy. Nature 458(7237):438–444.

15. Chen Y, Zhang Y, & Guo X (2017) Proteasome dysregulation in human cancer: implications for clinical therapies. Cancer Metastasis Rev 36(4):703–716.

16. Myeku N & Duff KE (2018) Targeting the 26S Proteasome To Protect Against Proteotoxic Diseases. Trends in molecular medicine 24(1):18–29.

17. Guo X, Huang X, & Chen MJ (2017) Reversible phosphorylation of the 26S proteasome. Protein & cell 8(4):255–272.

18. VerPlank JJS & Goldberg AL (2017) Regulating protein breakdown through proteasome phosphorylation. Biochem J 474(19):3355–3371.

19. Li D, et al. (2015) c-Abl regulates proteasome abundance by controlling the ubiquitin-proteasomal degradation of PSMA7 subunit. Cell Rep 10(4):484–496.

20. Liu X, et al. (2006) Interaction between c-Abl and Arg tyrosine kinases and proteasome subunit PSMA7 regulates proteasome degradation. Mol Cell 22(3):317–327.

21. Hunter T (2014) The genesis of tyrosine phosphorylation. Cold Spring Harbor perspectives in biology 6(5):a020644.

22. Kolch W & Pitt A (2010) Functional proteomics to dissect tyrosine kinase signalling pathways in cancer. Nat Rev Cancer 10(9):618–629.

23. Tibes R, Trent J, & Kurzrock R (2005) Tyrosine kinase inhibitors and the dawn of molecular cancer therapeutics. Annu Rev Pharmacol Toxicol 45:357–384.

24. Casaletto JB & McClatchey AI (2012) Spatial regulation of receptor tyrosine kinases in development and cancer. Nat Rev Cancer 12(6):387–400.

25. Sharma K, et al. (2014) Ultradeep human phosphoproteome reveals a distinct regulatory nature of Tyr and Ser/Thr-based signaling. Cell Rep 8(5):1583–1594.

26. Alonso A, et al. (2004) Protein tyrosine phosphatases in the human genome. Cell 117(6):699–711.

27. Sun H & Tonks NK (1994) The coordinated action of protein tyrosine phosphatases and kinases in cell signaling. Trends Biochem Sci 19(11):480–485.

28. Julien SG, Dubé N, Hardy S, & Tremblay ML (2011) Inside the human cancer tyrosine phosphatome. Nat Rev Cancer 11(1):35–49.

29. Frankson R, et al. (2017) Therapeutic Targeting of Oncogenic Tyrosine Phosphatases. Cancer Res 77(21):5701–5705.

30. Zhang ZY (2017) Drugging the Undruggable: Therapeutic Potential of Targeting Protein Tyrosine Phosphatases. Accounts of chemical research 50(1):122–129.

31. Thinon E, et al. (2014) Global profiling of co- and post-translationally N-myristoylated proteomes in human cells. Nature communications 5:4919.

32. Muppirala M, Gupta V, & Swarup G (2013) Emerging role of tyrosine phosphatase, TCPTP, in the organelles of the early secretory pathway. Biochimica et biophysica acta 1833(5):1125–1132.

33. Guerrero C, Milenkovic T, Przulj N, Kaiser P, & Huang L (2008) Characterization of the proteasome interaction network using a QTAX-based tag-team strategy and protein interaction network analysis. Proc Natl Acad Sci U S A 105(36):13333–13338.

34. Guo X, et al. (2016) Site-specific proteasome phosphorylation controls cell proliferation and tumorigenesis. Nat Cell Biol 18(2):202–212.

35. Huang X, Luan B, Wu J, & Shi Y (2016) An atomic structure of the human 26S proteasome. Nature structural & molecular biology 23(9):778–785.

36. Shibahara T, Kawasaki H, & Hirano H (2002) Identification of the 19S regulatory particle subunits from the rice 26S proteasome. Eur J Biochem 269(5):1474–1483.

37. Kimura Y, et al. (2003) N-Terminal modifications of the 19S regulatory particle subunits of the yeast proteasome. Arch Biochem Biophys 409(2):341–348.

38. Kimura A, Kato Y, & Hirano H (2012) N-myristoylation of the Rpt2 subunit regulates intracellular localization of the yeast 26S proteasome. Biochemistry 51(44):8856–8866.

39. Kimura A, Kurata Y, Nakabayashi J, Kagawa H, & Hirano H (2016) N-Myristoylation of the Rpt2 subunit of the yeast 26S proteasome is implicated in the subcellular compartment-specific protein quality control system. J Proteomics 130:33–41.

40. Zong C, et al. (2006) Regulation of murine cardiac 20S proteasomes: role of associating partners. Circ Res 99(4):372–380.

41. Wang X, et al. (2007) Mass spectrometric characterization of the affinity-purified human 26S proteasome complex. Biochemistry 46(11):3553–3565.

42. Kikuchi J, et al. (2010) Co- and post-translational modifications of the 26S proteasome in yeast. Proteomics 10(15):2769–2779.

43. Albert S, et al. (2017) Proteasomes tether to two distinct sites at the nuclear pore complex. Proc Natl Acad Sci U S A 114(52):13726–13731.

44. Ramachandran KV & Margolis SS (2017) A mammalian nervous-system-specific plasma membrane proteasome complex that modulates neuronal function. Nature structural & molecular biology.

45. Kohn AD, Summers SA, Birnbaum MJ, & Roth RA (1996) Expression of a constitutively active Akt Ser/Thr kinase in 3T3-L1 adipocytes stimulates glucose uptake and glucose transporter 4 translocation. J Biol Chem 271(49):31372–31378.

46. Moritz A, et al. (2010) Akt-RSK-S6 kinase signaling networks activated by oncogenic receptor tyrosine kinases. Science signaling 3(136):ra64.

47. Cross DA, et al. (2014) AZD9291, an irreversible EGFR TKI, overcomes T790M-mediated resistance to EGFR inhibitors in lung cancer. Cancer discovery 4(9):1046–1061.

48. Wang J & Gray NS (2015) SnapShot: Kinase Inhibitors II. Mol Cell 58(4):710.e711.

49. Kamps MP, Buss JE, & Sefton BM (1985) Mutation of NH2-terminal glycine of p60src prevents both myristoylation and morphological transformation. Proc Natl Acad Sci U S A 82(14):4625–4628.

50. Li X, et al. (1998) Generation of destabilized green fluorescent protein as a transcription reporter. J Biol Chem 273(52):34970–34975.

51. Hein MY, et al. (2015) A human interactome in three quantitative dimensions organized by stoichiometries and abundances. Cell 163(3):712–723.

52. St-Denis N, et al. (2016) Phenotypic and Interaction Profiling of the Human Phosphatases Identifies Diverse Mitotic Regulators. Cell Rep 17(9):2488–2501.

53. Kumar P, et al. (2017) A Human Tyrosine Phosphatase Interactome Mapped by Proteomic Profiling. J Proteome Res 16(8):2789–2801.

54. van Vliet C, et al. (2005) Selective regulation of tumor necrosis factor-induced Erk signaling by Src family kinases and the T cell protein tyrosine phosphatase. Nature immunology 6(3):253–260.

55. Zhang S, et al. (2009) Acquisition of a potent and selective TC-PTP inhibitor via a stepwise fluorophore-tagged combinatorial synthesis and screening strategy. Journal of the American Chemical Society 131(36):13072–13079.

56. Baselga J, et al. (2010) Phase I safety, pharmacokinetics, and inhibition of SRC activity study of saracatinib in patients with solid tumors. Clin Cancer Res 16(19):4876–4883.

57. Ye DZ & Field J (2012) PAK signaling in cancer. Cellular logistics 2(2):105–116.

58. Zhen Y, et al. (2015) Clinicopathologic significance of legumain overexpression in cancer: a systematic review and meta-analysis. Scientific reports 5:16599.

59. Simanshu DK, Nissley DV, & McCormick F (2017) RAS Proteins and Their Regulators in Human Disease. Cell 170(1):17–33.

60. Alevizopoulos K, Calogeropoulou T, Lang F, & Stournaras C (2014) Na+/K+ ATPase inhibitors in cancer. Current drug targets 15(10):988–1000.

61. Formisano L, et al. (2015) Src inhibitors act through different mechanisms in Non-Small Cell Lung Cancer models depending on EGFR and RAS mutational status. Oncotarget 6(28):26090–26103.

62. Tanaka K, et al. (1990) Molecular cloning of cDNA for proteasomes from rat liver: primary structure of component C3 with a possible tyrosine phosphorylation site. Biochemistry 29(15):3777–3785.

63. Hemmis CW, Heard SC, & Hill CP (2019) Phosphorylation of Tyr-950 in the proteasome scaffolding protein RPN2 modulates its interaction with the ubiquitin receptor RPN13. J Biol Chem 294(25):9659–9665.

64. Tang M, et al. (2017) Downregulation of SIRT7 by 5-fluorouracil induces radiosensitivity in human colorectal cancer. Theranostics 7(5):1346–1359.

65. Scott JD & Pawson T (2009) Cell signaling in space and time: where proteins come together and when they’re apart. Science 326(5957):1220–1224.

66. Kim YC & DeMartino GN (2011) C termini of proteasomal ATPases play nonequivalent roles in cellular assembly of mammalian 26 S proteasome. J Biol Chem 286(30):26652–26666.

67. Murata S, Takahama Y, Kasahara M, & Tanaka K (2018) The immunoproteasome and thymoproteasome: functions, evolution and human disease. Nature immunology 19(9):923–931.

68. Bose S, Brooks P, Mason GG, & Rivett AJ (2001) gamma-Interferon decreases the level of 26 S proteasomes and changes the pattern of phosphorylation. Biochem J 353(Pt 2):291–297.

69. Liu X, et al. (2020) Reversible phosphorylation of Rpn1 regulates 26S proteasome assembly and function. Proc Natl Acad Sci U S A 117(1):328–336.

70. Lokireddy S, Kukushkin NV, & Goldberg AL (2015) cAMP-induced phosphorylation of 26S proteasomes on Rpn6/PSMD11 enhances their activity and the degradation of misfolded proteins. Proc Natl Acad Sci U S A 112(52):E7176–7185.

71. Satoh K, Sasajima H, Nyoumura K-i, Yokosawa H, & Sawada H (2000) Assembly of the 26S Proteasome Is Regulated by Phosphorylation of the p45/Rpt6 ATPase Subunit. Biochemistry 40(2):314–319.

72. Manning G, Whyte DB, Martinez R, Hunter T, & Sudarsanam S (2002) The protein kinase complement of the human genome. Science 298(5600):1912–1934.

73. McCutchen-Maloney SL, et al. (2000) cDNA cloning, expression, and functional characterization of PI31, a proline-rich inhibitor of the proteasome. J Biol Chem 275(24):18557–18565.

74. Gorbea C, Goellner GM, Teter K, Holmes RK, & Rechsteiner M (2004) Characterization of mammalian Ecm29, a 26 S proteasome-associated protein that localizes to the nucleus and membrane vesicles. J Biol Chem 279(52):54849–54861.

75. Heun Y, et al. (2019) The Phosphatase SHP-2 Activates HIF-1α in Wounds In Vivo by Inhibition of 26S Proteasome Activity. International journal of molecular sciences 20(18).

76. Barr AJ (2010) Protein tyrosine phosphatases as drug targets: strategies and challenges of inhibitor development. Future medicinal chemistry 2(10):1563–1576.

77. Tiganis T, Bennett AM, Ravichandran KS, & Tonks NK (1998) Epidermal growth factor receptor and the adaptor protein p52Shc are specific substrates of T-cell protein tyrosine phosphatase. Mol Cell Biol 18(3):1622–1634.

78. Wei J, et al. (2019) Targeting REGNASE-1 programs long-lived effector T cells for cancer therapy. Nature 576(7787):471–476.

79. Manguso RT, et al. (2017) In vivo CRISPR screening identifies Ptpn2 as a cancer immunotherapy target. Nature 547(7664):413–418.

80. Wiede F, et al. (2020) PTPN2 phosphatase deletion in T cells promotes anti-tumour immunity and CAR T-cell efficacy in solid tumours. Embo j 39(2):e103637.

81. Feng Y, et al. (2017) Genetic variants of PTPN2 are associated with lung cancer risk: a re-analysis of eight GWASs in the TRICL-ILCCO consortium. Scientific reports 7(1):825.

82. Guo X, et al. (2016) Site-specific proteasome phosphorylation controls cell proliferation and tumorigenesis. Nat Cell Biol 18(2):202–212.

83. Kisselev AF & Goldberg AL (2005) Monitoring activity and inhibition of 26S proteasomes with fluorogenic peptide substrates. Methods Enzymol 398:364–378.

84. Guo X, et al. (2011) UBLCP1 is a 26S proteasome phosphatase that regulates nuclear proteasome activity. Proc Natl Acad Sci U S A 108(46):18649–18654.

85. Wiśniewski JR, Zougman A, Nagaraj N, & Mann M (2009) Universal sample preparation method for proteome analysis. Nature methods 6(5):359–362.

86. Cox J & Mann M (2008) MaxQuant enables high peptide identification rates, individualized p.p.b.-range mass accuracies and proteome-wide protein quantification. Nat Biotechnol 26(12):1367–1372.

87. Mi H, et al. (2019) Protocol Update for large-scale genome and gene function analysis with the PANTHER classification system (v.14.0). Nature protocols 14(3):703–721.

88. Benjamini Y, Hochberg, Y. (1995) Controlling the False Discovery Rate: A Practical and Powerful Approach to Multiple Testing. J. R. Stat. Soc. Ser. B. 57(1):289–300.

89. Szklarczyk D, et al. (2019) STRING v11: protein-protein association networks with increased coverage, supporting functional discovery in genome-wide experimental datasets. Nucleic acids research 47(D1):D607–d613.

